# Efficacy of the oral nucleoside prodrug GS-5245 (Obeldesivir) against SARS-CoV-2 and coronaviruses with pandemic potential

**DOI:** 10.1101/2023.06.27.546784

**Authors:** David R. Martinez, Fernando R. Moreira, Mark R. Zweigart, Kendra L. Gully, Gabriela De la Cruz, Ariane J. Brown, Lily E. Adams, Nicholas Catanzaro, Boyd Yount, Thomas J. Baric, Michael L. Mallory, Helen Conrad, Samantha R. May, Stephanie Dong, D. Trevor Scobey, Stephanie A. Montgomery, Jason Perry, Darius Babusis, Kimberly T. Barrett, Anh-Hoa Nguyen, Anh-Quan Nguyen, Rao Kalla, Roy Bannister, John P. Bilello, Joy Y. Feng, Tomas Cihlar, Ralph S. Baric, Richard L. Mackman, Alexandra Schäfer, Timothy P. Sheahan

## Abstract

Despite the wide availability of several safe and effective vaccines that can prevent severe COVID-19 disease, the emergence of SARS-CoV-2 variants of concern (VOC) that can partially evade vaccine immunity remains a global health concern. In addition, the emergence of highly mutated and neutralization-resistant SARS-CoV-2 VOCs such as BA.1 and BA.5 that can partially or fully evade (*1*) many therapeutic monoclonal antibodies in clinical use underlines the need for additional effective treatment strategies. Here, we characterize the antiviral activity of GS-5245, Obeldesivir (ODV), an oral prodrug of the parent nucleoside GS-441524, which targets the highly conserved RNA-dependent viral RNA polymerase (RdRp). Importantly, we show that GS-5245 is broadly potent in vitro against alphacoronavirus HCoV-NL63, severe acute respiratory syndrome coronavirus (SARS-CoV), SARS-CoV-related Bat-CoV RsSHC014, Middle East Respiratory Syndrome coronavirus (MERS-CoV), SARS-CoV-2 WA/1, and the highly transmissible SARS-CoV-2 BA.1 Omicron variant in vitro and highly effective as antiviral therapy in mouse models of SARS-CoV, SARS-CoV-2 (WA/1), MERS-CoV and Bat-CoV RsSHC014 pathogenesis. In all these models of divergent coronaviruses, we observed protection and/or significant reduction of disease metrics such as weight loss, lung viral replication, acute lung injury, and degradation in pulmonary function in GS-5245-treated mice compared to vehicle controls. Finally, we demonstrate that GS-5245 in combination with the main protease (M^pro^) inhibitor nirmatrelvir had increased efficacy in vivo against SARS-CoV-2 compared to each single agent. Altogether, our data supports the continuing clinical evaluation of GS-5245 in humans infected with COVID-19, including as part of a combination antiviral therapy, especially in populations with the most urgent need for more efficacious and durable interventions.

## INTRODUCTION

The emergence of three highly pathogenic novel coronaviruses (CoVs) into immunologically naïve human populations in the last two decades underlines an urgent need to develop broad-acting countermeasures. While broad-spectrum vaccines (*2–4*) and monoclonal antibodies (*5–8*) show promise in animal models against the sarbecovirus subgenus, the spike protein has extensively mutated throughout the COVID-19 pandemic and has partially and/or fully evaded vaccine and monoclonal antibody therapies in clinical use (*1*). In contrast, highly conserved viral enzymes, like the RNA-dependent RNA polymerase (RdRp, non-structural protein 12, NSP12) or main protease (i.e. M^pro^, NSP5) are more genetically stable and thus represent rational targets for broad-based antivirals. As broadly acting antivirals targeting the CoV RdRp, both remdesivir and molnupiravir have antiviral activity in vitro and in vivo against SARS-CoV-2 and divergent coronaviruses (*9–16*) and have been deployed for human use in the COVID-19 pandemic (*17, 18*). The M^pro^ inhibitor nirmatrelvir (PF-07321332 or PF-332), the active antiviral agent in Paxlovid™, exerts strong antiviral activity in vitro and in SARS-CoV-2 animal models (*19*). Importantly, Veklury® (remdesivir), Lagevrio™ (molnupiravir), and Paxlovid™ (nirmatrelivr/ritonavir) all improve outcomes in COVID-19-infected patients when given early in the course of infection (*18, 20–22*) and thus far have retained their antiviral activity against SARS-CoV-2 VOC including Omicron (*16, 23*). However, with the continued emergence of SARS-CoV-2 variants and increased use of antiviral monotherapy, it is critical to strengthen our armamentarium of orally bioavailable drugs and their combinations to treat COVID-19 in all populations, reduce its impact on the health care system, and minimize the development of antiviral resistance.

Here, we tested the antiviral efficacy of an oral prodrug, GS-5245 (ODV), of the nucleoside analog, GS-441524 (*24*). The prodrug is rapidly cleaved pre-systemically to generate GS-441524 into systemic circulation at higher exposures than what is achieved through direct oral dosing of GS-441524. We demonstrate that GS-5245 has broad therapeutic efficacy against endemic, enzootic, and pandemic coronaviruses in vitro and in vivo following oral delivery, including NL63, bat SARS-related RsSHC014-CoV, SARS-CoV, MERS-CoV, the ancestral SARS-CoV-2 WA1, and the highly transmissible SARS-CoV-2 Omicron variant. Moreover, therapy with a combination of GS-5245 and nirmatrelvir resulted in increased efficacy against SARS-CoV-2 replication in mice than either agent alone. These results support the continued exploration of GS-5245 in human clinical trials and confirms the need to develop an antiviral COVID-19 combination therapy in humans.

## RESULTS

### GS-5245 is broadly active against enzootic, endemic, and pandemic coronaviruses in primary human airway cells

We first evaluated the antiviral activity of the GS-5245, its parent nucleoside GS-441524, remdesivir, and the M^pro^ inhibitor PF-07321332 (PF-332), against a SARS-CoV-2 WA/1 nanoluciferase reporter virus in A549 cells that overexpress human ACE2 (Fig. 1A). SARS-CoV-2 was strongly inhibited by these antivirals with EC_50_ values of 0.74, 4.6, 0.19, and 0.07 µM for GS-5245, GS-441524, remdesivir, and PF-332, respectively, and where applicable, consistent with previously reported in vitro tests (*11, 19*). The improved potency for the prodrug GS-5245, compared to parent GS-441524, was also observed in other SARS-CoV-2 cell culture assessments and is thought to be due to the improved permeability properties, such as unexpectedly favorable interactions with intestinal nucleoside transporters, of the prodrug (*24*). We observed a strong reduction in SARS-CoV-2-expressed nanoluciferase activity (Fig. S1A) without cytotoxicity (Fig. S1B). To assess the antiviral breadth of GS-5245 against alphacoronavirus, we designed a nanoluciferase (nLuc)-expressing NL63 infectious clone and recovered recombinant reporter virus (Fig. S2A). NL63nLuc replicated with similar kinetics compared to a wild-type isolate (Fig. S2B) and expressed nanoluciferase to high levels in infected cells (Fig. S2C). Using NL63nLuc virus, we then tested the antiviral activity of GS-441524, remdesivir, and GS-5245 and observed robust antiviral activity in LLC-MK2 cells with respective EC_50_ values of 0.52, 0.49, and 0.62 µM (Fig. S3A and S3C) without evidence of cytotoxicity (Fig. S3B).

**Fig. 1.**
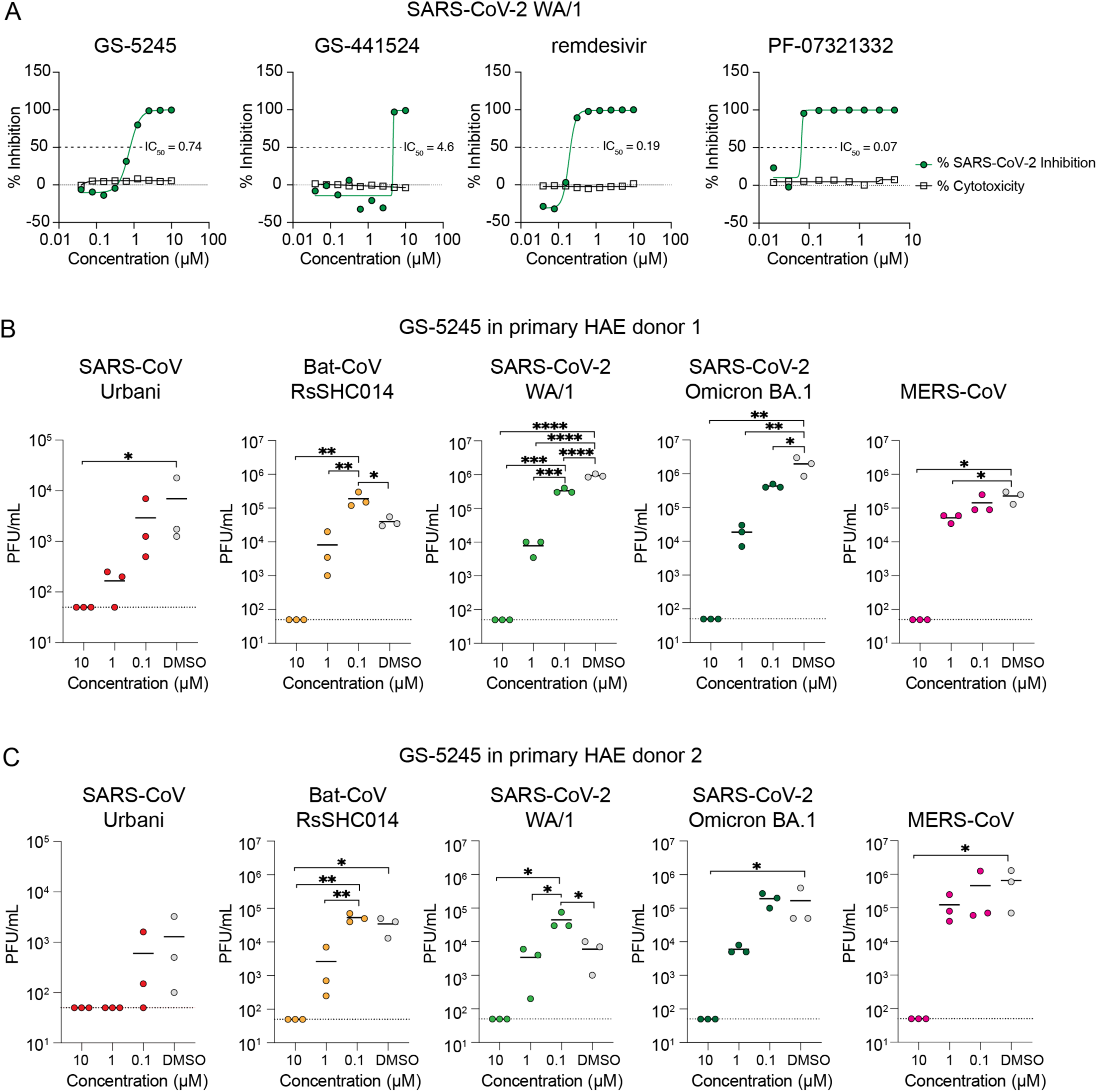
GS-5245 is broadly active against enzootic, endemic and pandemic coronaviruses in primary human airway cells. (A) The mean percent inhibition and cytotoxicity of SARS-CoV-2 replication by GS-5245 and control compounds GS-441524, remdesivir, and PF-07321332 in A459-hACE2 cells is shown (triplicate samples were analyzed). Data is representative of two independent experiments. (B) Antiviral activity of GS-5245 in primary human airway epithelial cell donor 1 against SARS-CoV and related bat-CoV SHC014, SARS-CoV-2 WA1 and SARS-CoV-2 Omicron BA.1 and MERS-CoV. Human airway epithelial cell cultures (HAE) were treated with different doses of drug or DMSO and then infected with SARS-CoV-2 at an MOI of 0.5. After 2 hours of infection, input virus was removed, cultures were washed once, and infectious virus titers in plaque forming units (PFU) was measured in apical washes after 72 hours. Cultures remained in the presence of drug for the duration of the study. (C) Antiviral activity of GS-5245 in primary human airway epithelial cell donor 2 against SARS-CoV and related bat-CoV SHC014, SARS-CoV-2 WA1, and SARS-CoV-2 Omicron BA.1 and MERS-CoV. This was performed similarly but independent of the study in panel B. For B and C, each symbol represents the virus titer per triplicate culture and the line is at the mean and * indicates p<0.05 as determined by One-Way ANOVA/Tukey’s multiple comparison test.

To further evaluate the breadth of GS-5245 antiviral activity against enzootic, endemic, epidemic, and pandemic coronaviruses, including highly transmissible SARS-CoV-2 variants, we evaluated GS-5245 in primary human airway epithelial (HAE) cells from two different human donors. The HAE platform is a highly biologically relevant three-dimensional culture system which models the structure and cellular complexity of the conducting airway and importantly contains epithelial target cells of the CoVs (*25*). Importantly, we observed a GS-5245 dose-dependent reduction in infectious virus production for all viruses tested including SARS-CoV Urbani, Bat-CoV RsSHC014, SARS-CoV-2 WA/1, the highly transmissible Omicron BA.1 variant, and MERS-CoV in HAE derived from two unique human donors (Figs. 1B and 1C). In addition, we did not observe measurable cytotoxicity of GS-5245 at the tested concentrations in HAE (Fig. S4). These studies demonstrate that the prodrug GS-5245 effectively releases parent GS-441524 in these cell experiments to generate the active triphosphate metabolite leading to antiviral activity. In conclusion, GS-441524 has potent activity against a broad array of genetically distinct CoVs including current SARS-CoV-2 VOC in the cell lines employed such as the biologically relevant primary HAE.

Residues in the nsp12 RdRp polymerase F-motif (V557 and A558) and B-motif (T687) help position the template in the active site (*26*). The conservation of residues in the active site that could impact incorporation, as well as residues responsible for both delayed chain termination and template-dependent inhibition. While the RdRp protein surface amino acid residue conservation was as low as 59% in HCoV-NL63 compared to SARS-CoV-2, the residues which would directly impact the efficacy of GS-5245 were 100% conserved across these and several other human and zoonotic coronaviruses (Fig. S5), demonstrating the potential for broad antiviral activity of small molecular inhibitors like GS-5245.

### Efficacy of GS-5245 and molnupiravir against SARS-CoV-2 in BALB/c mice

To determine the optimal therapeutic dose of GS-5245 in mice, we performed a therapeutic dose-ranging study in SARS-CoV-2 MA10-infected (1×10^4^ plaque forming units; PFU) BALB/c mice. We initiated therapy 12 hours post infection (hpi) with vehicle or 3, 10, or 30 mg/kg GS-5245 diluted in vehicle and mice were dosed orally twice daily (bis in die; BID) through 4 days post infection (dpi). The daily systemic exposure of GS-441524 following oral dosing at 30 mg/kg of GS-5245 across the different mice strains ranged from 81-108 µM.h and no intact GS-5245 prodrug was observed (Fig. S6B). This exposure of GS-441524 is consistent with the daily GS-441524 exposures achieved following oral administration of a tri-ester prodrug of GS-441524, GS-621763, dosed at 30 mg/kg BID in our earlier efficacy studies (*24, 27*).

We also included a cohort treated orally with molnupiravir at 100 mg/kg BID, a human equivalent dose determined based on the area under the curve (AUC) exposure. Overall, there was a distinct GS-5245 dose-dependent reduction in virus replication and pathogenesis (Figs. 2A-F). Mice treated with 3 mg/kg GS-5245 had measurable weight loss similar to vehicle, but only trended towards reductions in virus replication, macroscopic lung discoloration, and viral-induced pulmonary dysfunction. A higher GS-5245 dose of 10 mg/kg improved efficacy, as observed by significant protection from weight loss (Fig. 2A), lung virus titer, and gross lung pathology (i.e. lung discoloration). However, protection from pulmonary dysfunction and histological measures of acute lung injury (ALI) was not observed in the 10 mg/kg GS-5245 group. Mice treated with 30 mg/kg of GS-5245 were protected against weight loss (Fig. 2A) and had undetectable virus in lung tissue (Fig. 2B) similar to the 100 mg/kg molnupiravir-treated group. Macroscopic (Fig. 2C) and microscopic (Figs. 2E and F) measures of lung pathology and physiologic measures of lung function by whole body plethysmography (WBP) (Fig. 2D) were all significantly improved as compared to the vehicle control arm. These data demonstrate the strong dose-dependent relationship between the dose of GS-5245 and protection from SARS-CoV-2 disease. Moreover, GS-5245 affords similar protection at a lower dose (30 mg/kg) to that of molnupiravir (100 mg/kg) in mice.

**Fig. 2.**
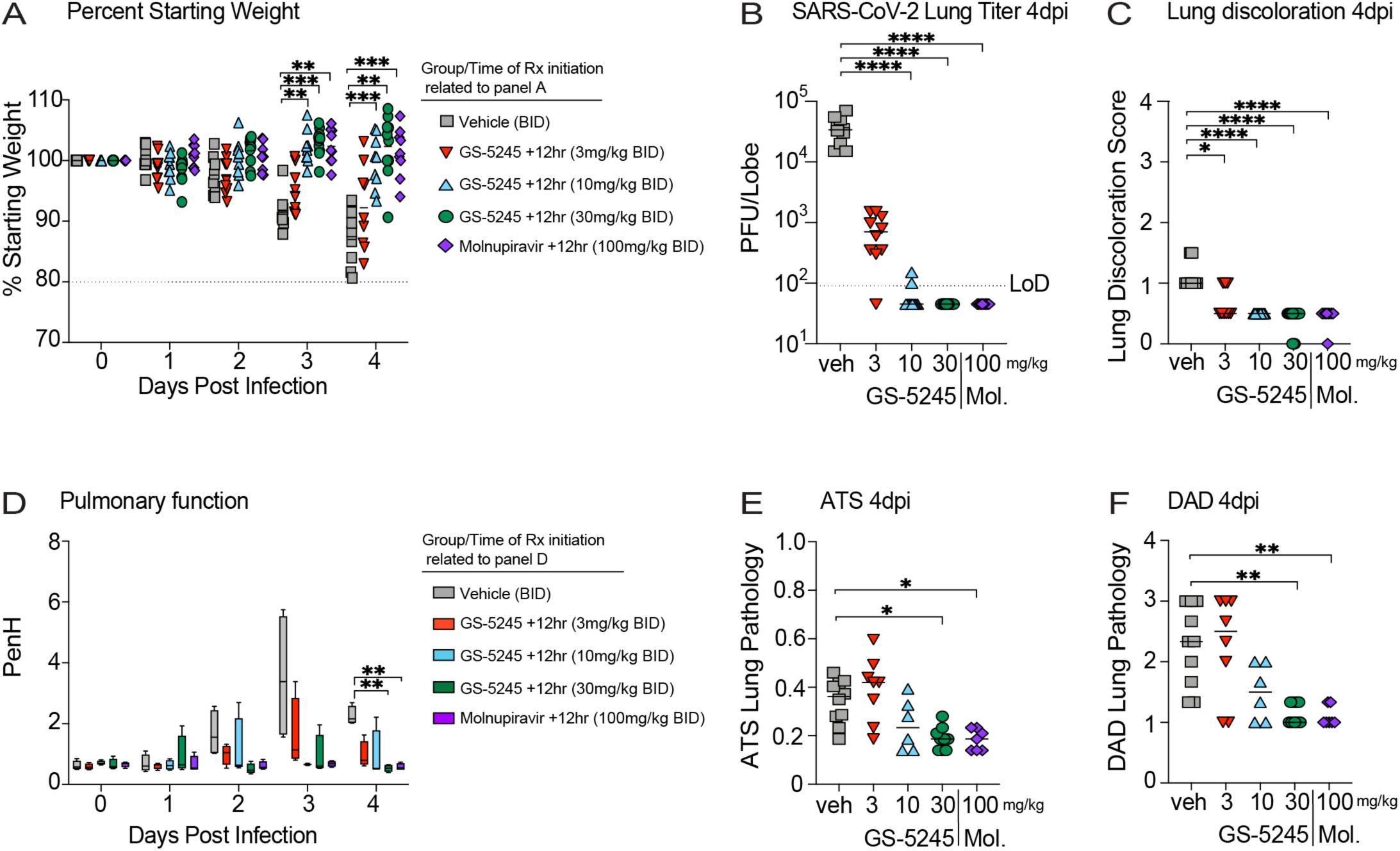
Dose-dependent therapeutic efficacy of GS-5245 against SARS-CoV-2 MA10 in BALB/c mice. (A) Percent starting weight through 4 dpi in 10-week-old female BALB/c mice infected with SARS-CoV-2 MA10 at 1 x 10^4^ PFU. Mice were treated BID with vehicle (veh: gray squares), 3 mg/kg GS-5245 (upside-down red triangles), 10 mg/kg GS-5245 (right-side-up blue triangles), 30 mg/kg GS-5245 (green circles), and 100 mg/kg molnupiravir (purple diamonds) starting at 12 hpi. N=10 mice per group. Rx = drug. Asterisks denote p values from a two-way ANOVA after a Dunnett’s multiple comparisons test. * p ≤ 0.05, ** p ≤ 0.01, *** p ≤ 0.001, **** p ≤ 0.0001 (B) SARS-CoV-2 MA10 lung infectious viral titers 4 dpi in mice treated with vehicle or GS-5245 at increasing concentrations and 100 mg/kg molnupiravir (Mol.). Asterisks denote p values from a Kruskal-Wallis test after a Dunnett’s multiple comparisons test. (C) Macroscopic lung discoloration at 4 dpi in therapeutically treated mice compared to vehicle. Asterisks denote p values from a Kruskal-Wallis test after a Dunnett’s multiple comparisons test. (D) Pulmonary function (PenH) monitored by whole-body plethysmography from day zero through 4 dpi in SARS-CoV-2-infected treated mice. Asterisks denote p values from a two-way ANOVA after a Dunnett’s multiple comparisons test. (E) Microscopic ATS acute lung injury pathology scoring at day 4 post infection in vehicle vs. GS-5245 or molnupiravir-treated mice. Asterisks denote p values from a Kruskal-Wallis test after a Dunnett’s multiple comparisons test. (F) Microscopic DAD acute lung injury pathology scoring at 4 dpi in vehicle, GS-5245, or molnupiravir-treated mice. Asterisks denote p values from a Kruskal-Wallis test after a Dunnett’s multiple comparisons test.

### Early treatment with GS-5245 reduces SARS-CoV-2 pathogenesis in BALB/c mice

To determine the time at which GS-5245 therapy fails to improve outcomes in mice, we performed a therapeutic efficacy study initiating treatment in SARS-CoV-2 MA10 infected (1×10^4^ PFU) BALB/c mice at 12, 24, and 36 hpi. We chose 30 mg/kg BID GS-5245 as it provided the most robust efficacy in prior studies. As expected, protection from virus replication and disease was dependent on the time of initiation of therapy (*14*). We observed complete protection from weight loss, lung viral replication, gross lung pathology, and degradation of pulmonary function (Fig. S7A-D) in the 12 and 24 hpi groups. In contrast, in the cohort where we initiated 30 mg/kg GS-5245 at 36 hpi, we did not observe protection from weight loss, lung pathology, or respiratory function, despite the significant decrease in lung viral replication compared to the vehicle group (Fig. S7A-D). Given the success of 30 mg/kg therapy at 24 hpi, we next aimed to determine the degree of protection with a lower dose of 10 mg/kg GS-5245 administered at the same timepoints post infection (Fig. S8). As we had seen with 30 mg/kg therapy, mice treated with 10 mg/kg GS-5245 at 12 hpi were protected from weight loss (Fig. S8A); had reduced virus lung titers (Fig. S8B), macroscopic lung discoloration (Fig. S8C), histologic acute lung injury scores (Fig. S8D, S8E), and degradation of pulmonary function (Fig. S8F). Treatment with 10 mg/kg at 24 or 36 hpi did not prevent body weight loss (Fig. S8A), but did reduce virus titer (Fig. S8B), gross pathology (Fig. S8C) and one of two histologic acute lung injury scores (Fig. S8D, 24 hpi only). Altogether, these data suggests that the therapeutic activity of GS-5245 at either 10 or 30 mg/kg is most effective early after infection (12 or 24 hpi) prior to the peak of viral replication, which is at 48 hpi in this SARS-CoV-2 mouse model (*28*).

### GS-5245 is effective against SARS-related Bat-CoV RsSHC014 in K18-hACE mice

We next aimed to understand the breadth of antiviral efficacy in vivo against coronaviruses that are more distantly related to SARS-CoV-2, including SARS-related Bat-CoV RsSHC014 and MERS-CoV, which frequently emerges. As we aimed to use transgenic mice on a different genetic background, C57BL/6, than that used in the above efficacy studies, we first performed a pharmacokinetic (PK) study with GS-5245 in K18-human angiotensin-convertase enzyme 2 (hACE2) mice and those with humanized human dipeptidyl peptidase 4 (hDPP4) (Fig. S6A), which are susceptible to RsSHC014 and MERS-CoV, respectively. After oral dosing with GS-5245 at 30 mg/kg, we measured the nucleoside parent GS-441524 in mouse plasma over time. We observed no differences in the plasma levels of GS-441524 in either transgenic line when compared to control BALB/c, supporting the use of similar oral doses of GS-5245 in our planned in vivo efficacy studies in these transgenic mice (Fig. S6B).

The SARS-like Bat-CoV RsSHC014 can efficiently replicate in human airway epithelial cells and can evade existing SARS-CoV-2 vaccines and mAb countermeasures against SARS-CoV (*4, 29*). Thus, we evaluated the therapeutic efficacy of GS-5245 in RsSHC014 infected K18-hACE2 mice initiating treatment at times post-infection (i.e. 12 and 24 hpi) with demonstrable success in SARS-CoV-2 models described above. Unlike the SARS-CoV-2 MA10 BALB/c model, marked weight loss is not a hallmark of RsSHC014 infection in K18-hACE2 mice through day 4, although we did observe increased weight in the infected animals dosed with 30 mg/kg initiated at 12 hpi compared to vehicle-treated mice (Fig. 3A). While we observed a trend in reduced virus replication (Fig. 3B) and gross lung pathology (Fig. 3C) with the 10 mg/kg dose, only the 30 mg/kg dose initiated at either 12 or 24 hpi significantly reduced virus lung titers, and gross lung pathology. As done in the SARS-CoV-2 MA10 model, we next evaluated histologic manifestations of acute lung injury (ALI) using two different scoring tools (Figs. 3D and E). Mice dosed with 30 mg/kg initiated at either 12 or 24 hpi had significant reductions in ALI. Thus, a significant reduction in SARS-related enzootic virus replication and disease was observed with GS-5245 therapy.

**Fig. 3.**
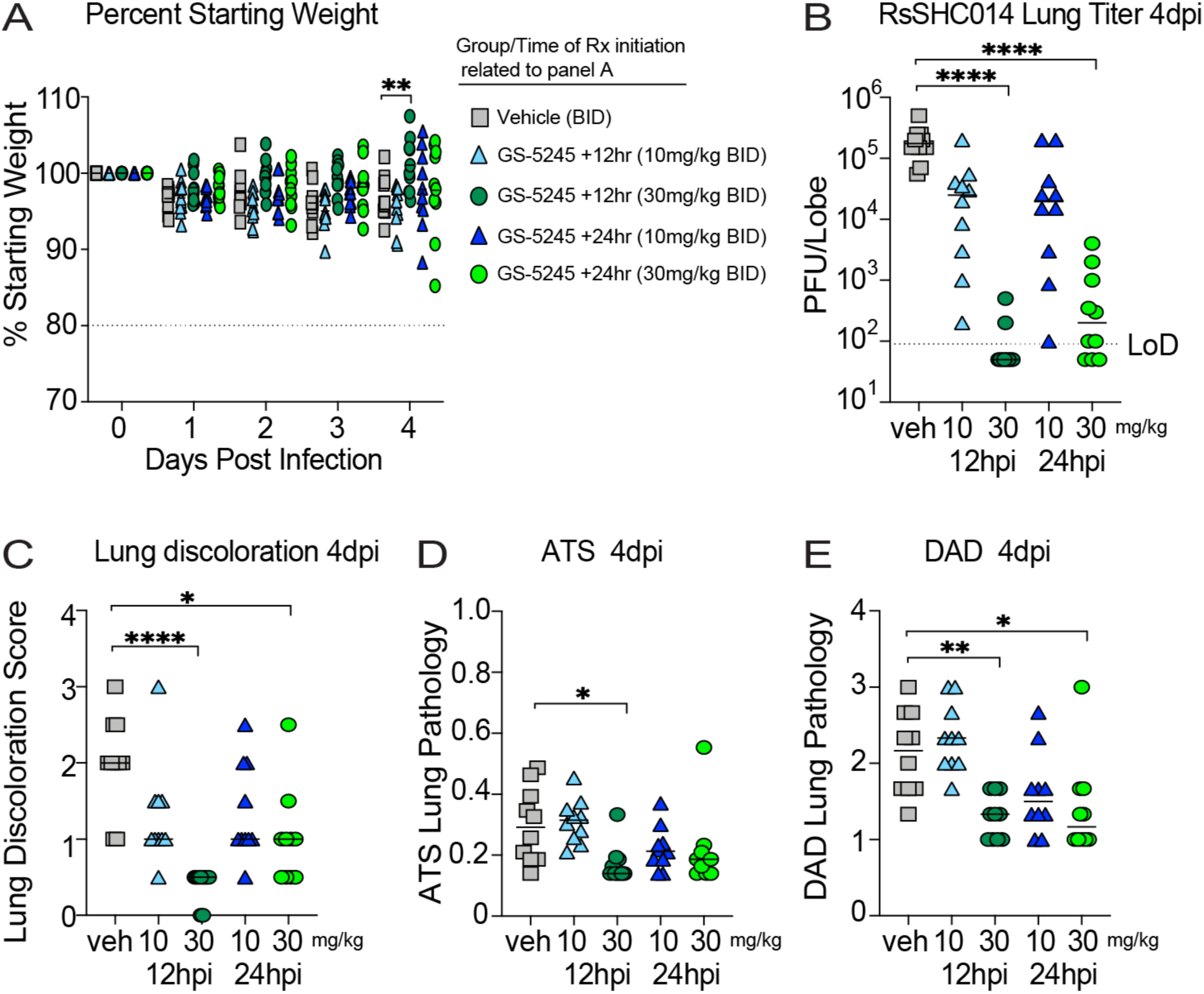
Therapeutic efficacy of GS-5245 against pre-emergent bat SARS-related RsSHC014-CoV in K18-hACE2 mice. (A) Percent starting weight in 10-week-old female K18-hACE2 mice infected with 1 x 10^4^ PFU therapeutically treated BID with vehicle (veh: gray squares), 10 mg/kg GS-5245 (right-side-up light blue triangles), 30 mg/kg GS-5245 (dark green circles) at 12 hpi, and 10 mg/kg GS-5245 (right-side-up dark blue triangles), and 30 mg/kg GS-5245 (light green circles) at 24 hpi. N=10 mice per group. Asterisks denote p values from a two-way ANOVA after a Dunnett’s multiple comparisons test. (B) RsSHC014-CoV lung infectious viral titers 4 dpi in mice treated with vehicle or GS-5245 at 10 and 30 mg/kg at either 12 or 24 hpi. Asterisks denote p values from a Kruskal-Wallis test after a Dunnett’s multiple comparisons test. (C) Macroscopic lung discoloration at 4 dpi in therapeutically treated K18-hACE2 mice compared to vehicle. Asterisks denote p values from a Kruskal-Wallis test after a Dunnett’s multiple comparisons test. (D) Microscopic ATS acute lung injury pathology scoring at day 4 post infection in vehicle vs. GS-5245-treated mice. Asterisks denote p values from a Kruskal-Wallis test after a Dunnett’s multiple comparisons test. (E) Microscopic DAD acute lung injury pathology scoring at day 4 post infection in vehicle vs. GS-5245-treated mice. Asterisks denote p values from a Kruskal-Wallis test after a Dunnett’s multiple comparisons test.

### The therapeutic efficacy of GS-5245 against SARS-CoV in BALB/c mice

With ∼10% mortality rate in humans, the highly virulent SARS-CoV strain emerged in 2002-2003 in Guangdong Province China ultimately causing over 8000 cases in 29 countries and over 800 deaths (*30, 31*). Therefore, we next aimed to evaluate the antiviral activity of GS-5245 against emerging coronaviruses distinct from SARS-CoV-2 with clear human epidemic potential. To increase the stringency of our in vivo assessment of GS-5245, we designed a therapeutic efficacy study using the highly pathogenic mouse-adapted SARS-CoV MA15 virus in BALB/c mice (*32*). Only early therapeutic intervention with 30 mg/kg initiated at 12 hpi protected from significant body weight loss (Fig. 4A). We observed a trend towards reduced virus replication (Fig. 4B), and gross lung pathology (Fig. 4C) with GS-5245 at 10 mg/kg, and significant reductions in these metrics was afforded by 30 mg/kg initiated at 12 hpi. As SARS-CoV MA15 infection causes mortality in BABL/c mice, we observed a high degree of protection in the 12 hpi 30 mg/kg group with 100% survival in this group compared to vehicle-treated mice which exhibited 80% mortality by day 4 post infection (Fig. 4D). We also observed 30% mortality in the 12 hpi 10mg/kg group. In contrast, both the 24 hpi 10 and 30 mg/kg treatments had similar mortality rates to the vehicle group. In our assessment of ALI using the ATS (Fig. 4E) or DAD (Fig. 4F) histologic scoring tools, only 30 mg/kg initiated at 12 hpi significantly reduced microscopic lung pathology. Although the lower dose of GS-5245 did not afford much protection in the metrics above, all dose groups had improved pulmonary function by 3 dpi (Fig. 4G). In addition, 30 mg/kg initiated at 12 hpi prevented the loss of pulmonary function observed in the vehicle on days 1 and 2 (Fig. 4G). Thus, early treatment with a 30 mg/kg dose of GS-5245 is highly protective against SARS-CoV disease symptoms and mortality in mice.

**Fig. 4.**
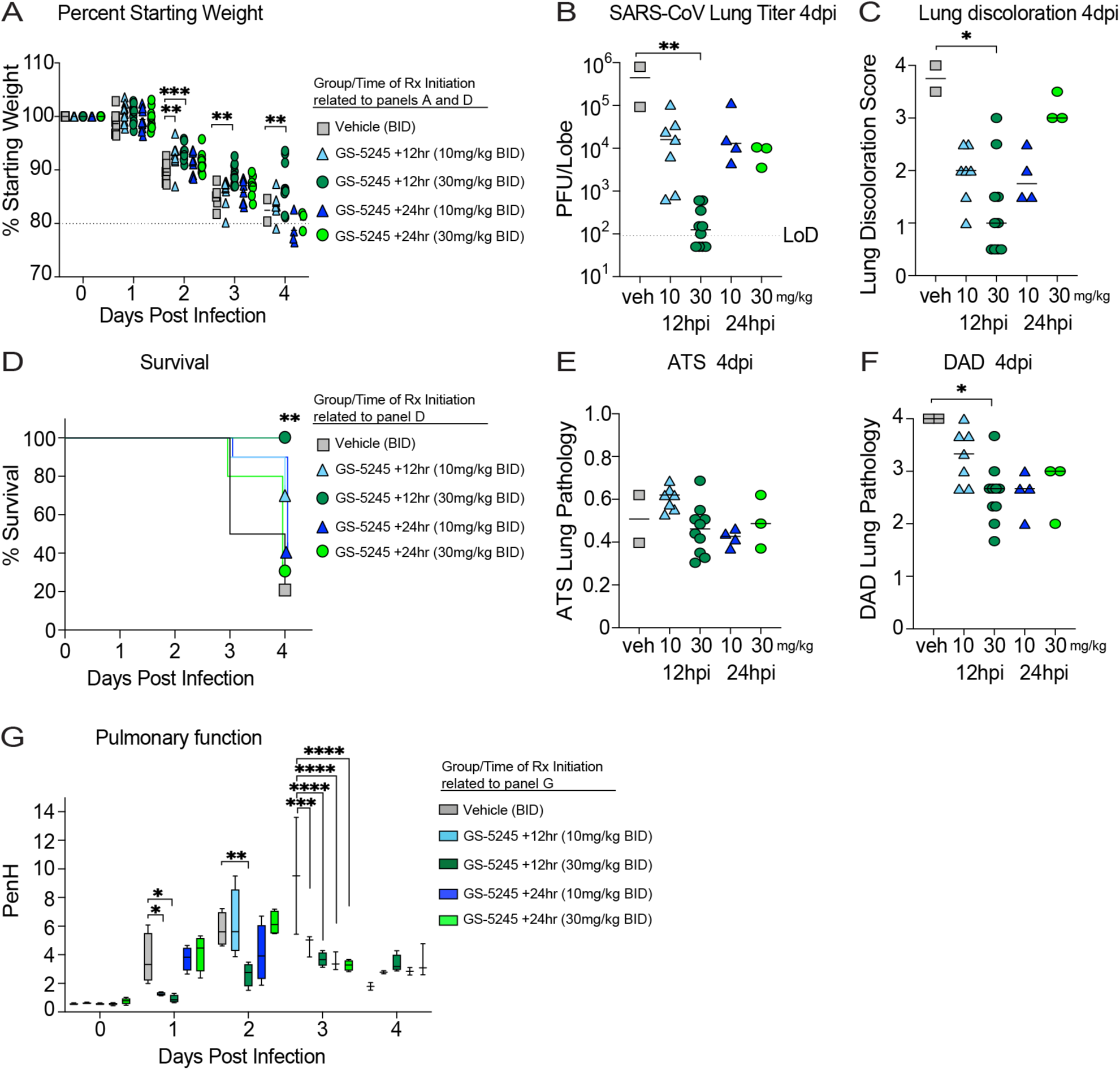
GS-5245 protects against SARS-CoV MA15 mortality in BALB/c mice. (A) Percent starting weight in 10-week-old female BALB/c mice infected with 1 x 10^4^ PFU therapeutically treated BID with vehicle (veh: gray squares), 10 mg/kg GS-5245 (right-side-up light blue triangles), 30 mg/kg GS-5245 (dark green circles) at 12 hpi, and 10 mg/kg GS-5245 (right-side-up dark blue triangles), and 30 mg/kg GS-5245 (light green circles) at 24 hpi. N=10 mice per group. Asterisks denote p values from a two-way ANOVA after a Dunnett’s multiple comparisons test. (B) SARS-CoV MA15 lung infectious viral titers 4 dpi in mice treated with vehicle or GS-5245 at 10 and 30 mg/kg at either 12 or 24 hpi. Asterisks denote p values from a Kruskal-Wallis test after a Dunnett’s multiple comparisons test. (C) Macroscopic lung discoloration at 4 dpi in therapeutically treated BALB/c mice compared to vehicle. Asterisks denote p values from a Kruskal-Wallis test after a Dunnett’s multiple comparisons test. (D) Percent survival in vehicle vs GS-5245-treated mice. Asterisks denote p values from a Log-rank Mantel-Cox test. (E) Microscopic ATS acute lung injury pathology scoring at 4 dpi in vehicle vs. GS-5245-treated mice. Asterisks denote p values from a Kruskal-Wallis test after a Dunnett’s multiple comparisons test. (F) Microscopic DAD acute lung injury pathology scoring at 4 dpi in vehicle vs. GS-5245-treated mice. Asterisks denote p values from a Kruskal-Wallis test after a Dunnett’s multiple comparisons test. (G) Pulmonary function monitored by whole-body plethysmography from day zero through 4 dpi in SARS-CoV-infected treated mice. Asterisks denote p values from a two-way ANOVA after a Dunnett’s multiple comparisons test.

### Successful in vivo efficacy with GS-5245 against MERS-CoV in DPP4 288/330-modified mice

We next aimed to assess therapeutic efficacy against MERS-CoV using a mouse model that utilizes a modified dipeptidyl peptidase 4 (DPP4) at amino acid positions 288 and 330 (*33, 34*). Since initiating therapy at 12 hpi offered the most protection in the SARS-CoV model above, we evaluated the therapeutic efficacy of GS-5245 at 10 or 30 mg/kg in MERS-CoV infected animals at 12 hpi. In agreement with the SARS-CoV-2, and SARS-CoV in vivo data above, GS-5245 at 30 mg/kg provided the strongest protection against signs of clinical disease including weight loss, lung viral titers, acute lung injury, and degradation in respiratory function (Figs. 5A-F). While the lower 10 mg/kg dose only afforded partial protection from weight loss (Fig. 5A), it markedly reduced lung viral replication (Fig. 5B). Unlike the 30 mg/kg dose, the lower 10 mg/kg dose did not reduce gross lung pathology (Fig. 5C), pulmonary function as measured by whole body plethysmography (Fig. 5D), or ALI as measured by quantitative histologic scoring tools (Figs. 5E and F). Thus, GS-5245 protects against MERS-CoV pathogenesis in mice when therapy is initiated soon after infection.

**Fig. 5.**
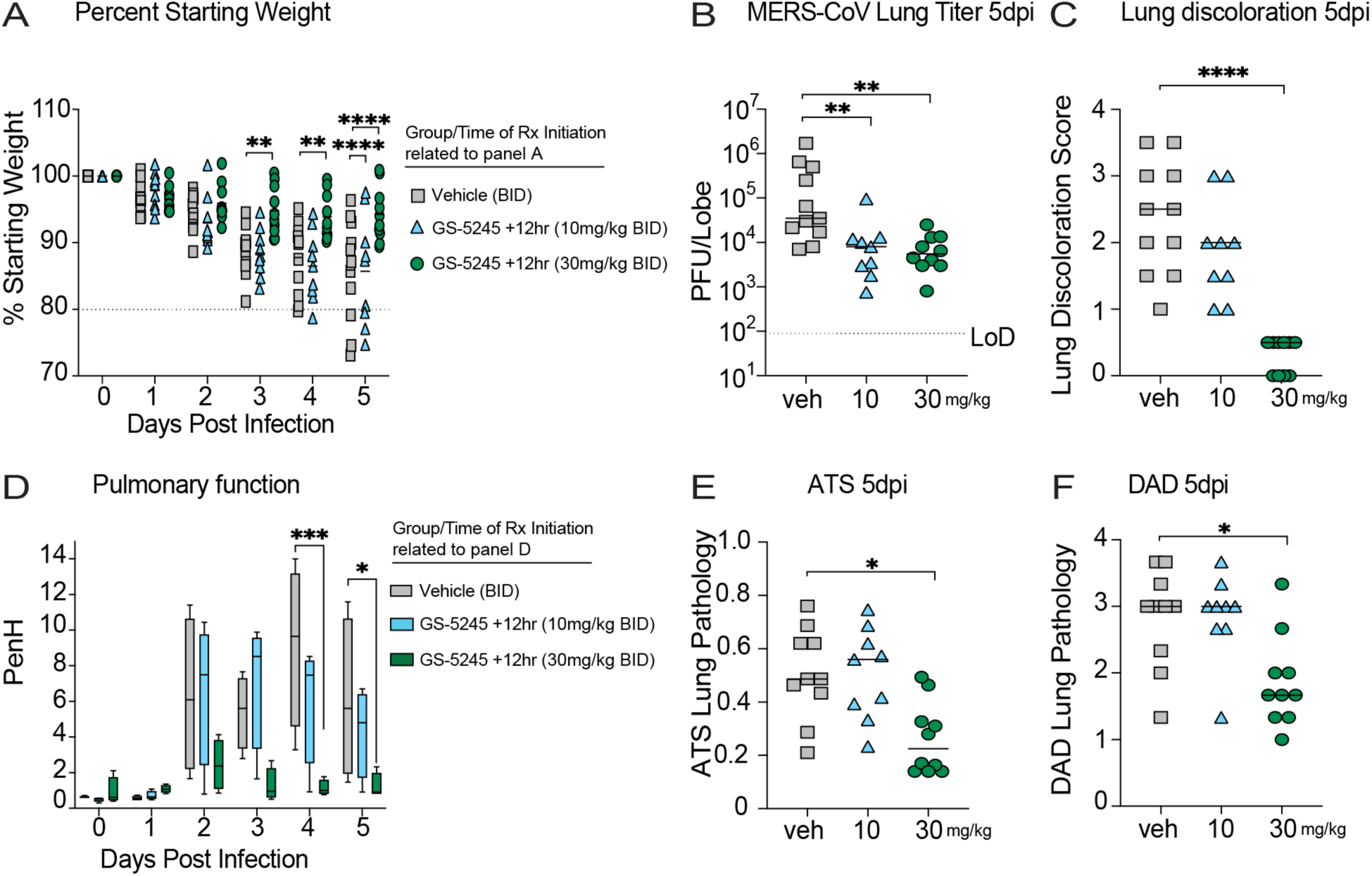
GS-5245 inhibits MERS-CoV disease in 288/330^++^ DPP4-modified mice. (A) Percent starting weight in 20-week-old male and female 288/330^++^ DPP4-modified mice infected with a mouse-adapted MERS-CoV with 5 x 10^4^ PFU. Mice were therapeutically treated BID starting at 12 hpi with vehicle (veh: gray squares), 10 mg/kg GS-5245 (right-side-up light blue triangles), 30 mg/kg GS-5245 (dark green circles). N=8 total mice per group. Male and female mice were distributed as equally as possible in each group. Asterisks denote p values from a two-way ANOVA after a Dunnett’s multiple comparisons test. (B) MERS-CoV lung infectious viral titers 5 dpi in mice treated with vehicle or GS-5245 at 10 and 30 mg/kg at 12 hpi. Asterisks denote p values from a Kruskal-Wallis test after a Dunnett’s multiple comparisons test. (C) Macroscopic lung discoloration at 5 dpi in therapeutically treated mice compared to vehicle. Asterisks denote p values from a Kruskal-Wallis test after a Dunnett’s multiple comparisons test. (D) Pulmonary function monitored by whole-body plethysmography from day zero through 5 dpi in MERS-CoV-infected treated 288/330^++^ DPP4 mice. Asterisks denote p values from a two-way ANOVA after a Dunnett’s multiple comparisons test. (E) Microscopic ATS acute lung injury pathology scoring at 5 dpi in vehicle vs. GS-5245-treated mice. Asterisks denote p values from a Kruskal-Wallis test after a Dunnett’s multiple comparisons test. (F) Microscopic DAD acute lung injury pathology scoring at 5 dpi in vehicle vs. GS-5245-treated mice. Asterisks denote p values from a Kruskal-Wallis test after a Dunnett’s multiple comparisons test.

### GS-5245 therapy is highly effective at reducing SARS-CoV-2 Omicron replication in K18-hACE2 mice

As we observed a high degree of protection against bat RsSHC014-CoV, SARS-CoV, MERS-CoV, and a mouse-adapted SARS-CoV-2 based on the Wuhan-1 isolate, we sought to evaluate GS-5245 against the highly transmissible Omicron (B.1.1.529/BA.1.) variant in K18-hACE2 mice. As previous studies did not find that B.1.1.529 caused severe disease in K18-hACE2 mice including weight loss or lung pathology (*35*), we performed a therapeutic efficacy study where the main readout was BA.1 replication in the lung. Consistent with Halfman *et al.,* we did not observe weight loss through day 4 post infection nor severe lung pathology in vehicle-treated mice (Figs. 6A and 6C-E). Mice treated with GS-5245 at 30 mg/kg 12 hpi had significantly lower lung viral titers relative to vehicle-treated groups at 2-, 3-, and 4-days post infection (Fig. 6B), and protection from macroscopic and microscopic lung pathology compared to vehicle (Figs. 6C, 6D, and 6E). We conclude that GS-5245 demonstrates a high degree of protection against lung viral replication in vivo against the highly transmissible BA.1 variant.

**Fig. 6.**
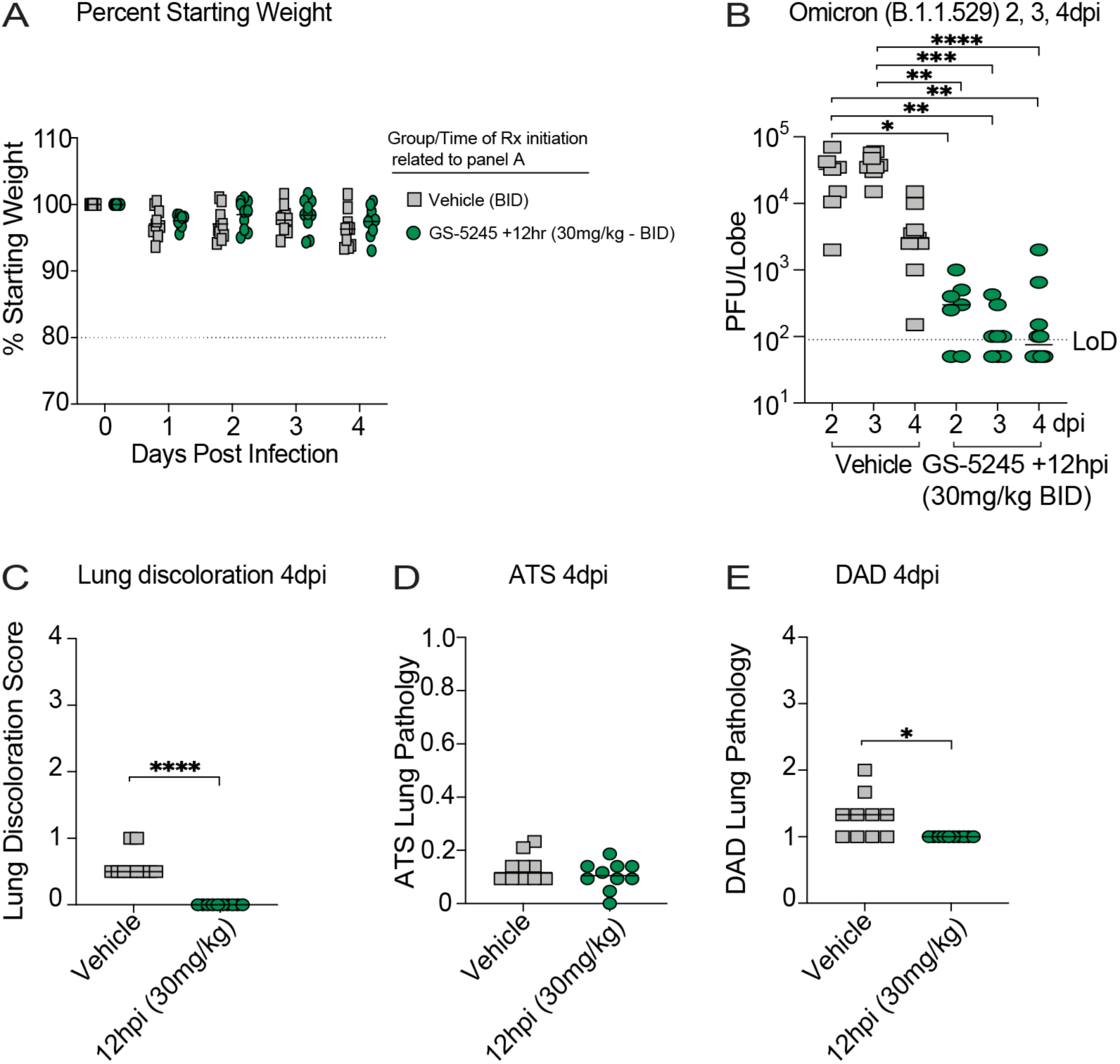
Therapeutic efficacy of GS-5245 against the highly transmissible and neutralization-resistant BA.1 (Omicron B.1.1.529) variant in K18-hACE2 mice. (A) Percent starting weight in 10-week-old female K18-hACE2 mice infected at 1 x 10^4^ PFU with a BA.1 clinical isolate. Mice were therapeutically treated BID starting at 12 hpi with vehicle (veh: gray squares) and 30 mg/kg GS-5245 (dark green circles). N=10 mice per group. Asterisks denote p values from a two-way ANOVA after a Dunnett’s multiple comparisons test. (B) BA.1 lung infectious viral titers 2, 3, and 4 dpi in mice treated with vehicle or GS-5245 and 30 mg/kg at 12 hpi. Asterisks denote p values from a Kruskal-Wallis test after a Dunnett’s multiple comparisons test. (C) Macroscopic lung discoloration at 4 dpi in therapeutically treated mice compared to vehicle. Asterisks denote p values from a Mann-Whitney test. (D) Microscopic ATS acute lung injury pathology scoring at 4 dpi in vehicle vs. GS-5245-treated mice. Asterisks denote p values from a Mann-Whitney test. (E) Microscopic DAD acute lung injury pathology scoring at 4 dpi in vehicle vs. GS-5245-treated mice. Asterisks denote p values from a Mann-Whitney test.

### Combination therapy of PF-07321332 and GS-5245 against SARS-CoV-2 in BALB/c mice

As the oral antiviral nirmatrelvir (PF-332) demonstrated a high degree of efficacy in mice and humans (*19, 20*), we sought to evaluate if combination of PF-332 with GS-5245 could further diminish SARS-CoV-2 pathogenesis in mice. PF-332 inhibits M^pro^ by preventing processing of the viral polyprotein whereas GS-5245 inhibits the RdRp by terminating transcription and replication (*36*) and/or excessive polymerase pausing (*37*). We first performed a dose-de-escalation experiment to determine the doses of PF-332 that provides optimal and suboptimal protection from SARS-CoV-2 replication and disease in mice. At 12 hpi, we initiated therapy with 400, 120, 40, or 12 mg/kg PF-332 BID or 1.2 mg/kg GS-5245 BID. We observed that PF-332 protected mice from weight loss (Fig. 7A), lung viral replication (Fig. 7B), lung discoloration (Fig. 7C), and degradation in pulmonary function (Fig. 7D) at the highest drug treatment doses but not at lower doses. Similarly, the 1.2 mg/kg GS-5245 dose was highly suboptimal and little protection was observed.

**Fig. 7.**
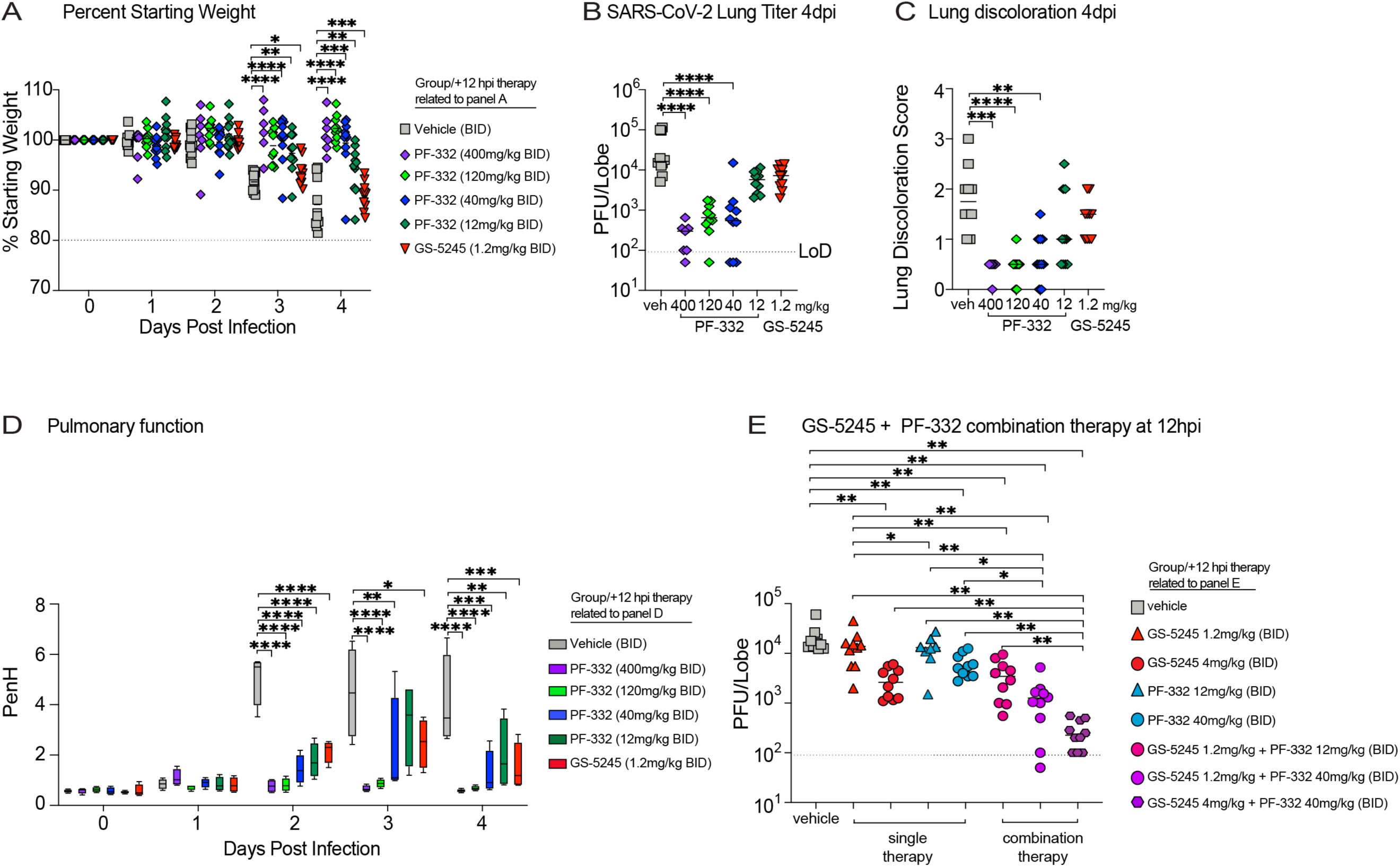
The effect of GS-5245 and PF-332 combination therapy on SARS-CoV-2 MA10 pathogenesis in BALB/c mice. (A) Percent starting weight in 10-week-old female BALB/c mice infected at 1 x 10^4^ PFU with a SARS-CoV-2 MA10. Mice were treated BID starting at 12 hpi with vehicle (veh: gray squares) 400 mg/kg PF-332 (purple diamonds), 120 mg/kg PF-332 (light green diamonds), 40 mg/kg PF-332 (blue diamonds), 12 mg/kg PF-332 (dark green diamonds), and 1.2 mg/kg GS-5245 (upside-down red triangles). N=10 mice per group. Asterisks denote p values from a two-way ANOVA after a Dunnett’s multiple comparisons test. (B) SARS-CoV-2 MA10 lung infectious viral titers 4 dpi in mice treated at 12 hpi with vehicle and at decreasing doses of PF-332 and a sub-protective dose of 1.2 mg/kg GS-5245. Asterisks denote p values from a Kruskal-Wallis test with a Dwass-Steel-Critchlow-Fligner procedure for pairwise comparisons. (C) Macroscopic lung discoloration at 4 dpi in therapeutically treated mice compared to vehicle. Asterisks denote p values from a Kruskal-Wallis test with a Dwass-Steel-Critchlow-Fligner procedure for pairwise comparisons. (D) Pulmonary function as measured by whole-body plethysmography in SARS-CoV-2-infected mice starting at day 0 through 4 dpi. (E) Sub-protective oral drug combination therapy and its effect on lung infectious SARS-CoV-2 viral titers. Vehicle (gray squares), 1.2 mg/kg GS-5245 (right-side-up red triangles), 4 mg/kg GS-5245 (red circles), 12 mg/kg PF-332 (right-side-up blue triangles), 40 mg/kg PF-332 (blue circles), combination 1.2 mg/kg GS-5245/12 mg/kg PF-332 (pink circles), combination 1.2 mg/kg GS-5245/40 mg/kg PF-332 (purple circles), combination 4 mg/kg GS-5245/40 mg/kg PF-332 (purple hexagons). Kruskal-Wallis test was used to compare the group samples, and Dwass-Steel-Critchlow-Fligner procedure was used for pairwise comparisons.

As virus replication is the main driver of disease in this model, we sought to determine if combination therapy initiated at 12 hpi with sub-optimal doses of PF-332 and GS-5245 would result in an increased reduction in virus lung titers than either drug alone. Thus, we designed a therapeutic efficacy study in mice infected with SARS-CoV-2 treated with suboptimal doses of single agents administered BID (GS-5245 at 1.2 or 4 mg/kg, PF-332 at 12 or 40 mg/kg) or several combinations of the two administered BID: “low dose combination” of 1.2 mg/kg GS-5245 + 12 mg/kg PF-332, “medium dose combination” of 1.2 mg/kg GS-5245 + 40 mg/kg PF-332, or “high dose combination” of 4 mg/kg GS-5245 + 40 mg/kg PF-332. As compared to vehicle-treated animals, SARS-CoV-2 lung titers were not reduced following therapy with suboptimal doses of single agents at multiple dose levels including 1.2 mg/kg of GS-5245 or PF-332 at 12 or 40 mg/kg (Fig. 7E). In contrast, intervention of GS-5245 at 4 mg/kg singly significantly reduced viral titers in the lung (Fig. 7E). Impressively, combination of GS-5245 at 1.2 mg/kg and PF-332 at 40 mg/kg resulted in significantly more profound levels of lung viral replication compared to vehicle and single agent groups. Similarly, by increasing the concentration of each component of combination therapy (4 mg/kg GS-5245 and 40 mg/kg PF-332), we observed a marked reduction in viral titers in the lung that was significantly lower than either single agent (Fig. 7E). Altogether, these data suggest that combination therapy of GS-5245 and PF-332 is highly effective at suppressing lung viral replication in mice even when combined at suboptimal doses.

## DISCUSSION

The SARS-CoV-2 spike protein, which is the primary target of neutralizing antibodies, has undergone extensive changes conferring an increased ability to evade existing antibody-based countermeasures (*1*). The emergence of the Omicron (BA.1) and several Omicron sub-lineages have continually eroded the neutralizing activity of vaccine-elicited antibodies in the COVID-19 vaccines (*38*). The natural waning of vaccine-elicited serum immune responses may contribute to the surge in infections with highly transmissible SARS-CoV-2 VOCs (*39*). Moreover, the emergence of more immune-evasive variants such as BQ.1.1 and XBB evade all human monoclonal antibodies in clinical use whereas antivirals like remdesivir remain active against these variants (*40*). In the setting of continued SARS-CoV-2 spike protein evolution and waning vaccine immunity, it is critical to develop orally bioavailable drugs that can broadly inhibit SARS-CoV-2 and its current and future VOCs.

Intravenously administered remdesivir (Veklury^®^) has received full Food and Drug Administration (FDA) approval; and orally administered molnupiravir and nirmatrelvir have received Emergency Use Authorization (EUA) for the treatment of COVID-19 in the U.S. Molnupiravir similarly exhibits broad-spectrum activity against zoonotic and pandemic coronaviruses in primary HAE cells, mice, and humans (*13, 21*). In contrast to remdesivir and molnupiravir, which target the RdRp albeit through different mechanisms of action, nirmatrelvir targets M^pro^. Nirmatrelvir alone showed a high degree of protection against SARS-CoV-2 pathogenesis in mice (*19*), and when combined with ritonavir (Paxlovid™) showed protection in human clinical trials (*20*). When given early after infection, remdesivir demonstrates efficacy and reduces risk of hospitalization due to COVID-19 (*22*). Consistent with this, early remdesivir treatment of SARS-CoV and SARS-CoV-2-infected mice offered the most protection against viral pathogenesis (*11, 14*).

Despite the promising efficacy of remdesivir, (*22*) its intravenous administration has been a major barrier for broad use due to the requirement of trained medical personnel to administer drug in outpatient settings such as infusion centers. To overcome this challenge, an oral prodrug (GS-5245; ODV) of the parent nucleoside GS-441524, was developed (*24*). GS-5245 ultimately forms the same active triphosphate in lung tissue as remdesivir. Here, we show that GS-5245 is highly effective at diminishing replication and disease pathogenesis for enzootic, epidemic, emerging, and pandemic coronaviruses including the highly transmissible SARS-CoV-2 Omicron variant following oral administration. The twice daily 30 mg/kg dose was effective across the different SARS-CoV-2 models and corresponded to daily GS-441524 exposures ranging from 81-108 µM.h. This exposure is consistent with the estimated GS-441524 exposure at efficacious oral doses of GS-5245 in a non-human primate SARS-CoV-2 infection model and the targeted exposures in humans (*24*). Thus, GS-5245 is potentially an additional oral antiviral for outpatients with COVID-19. In the context of pandemic preparedness, drugs targeting highly conserved viral proteins, like the RdRp, are advantageous as they may retain activity against future VOCs and emerging coronavirus threats. Viruses like SARS-CoV, MERS-CoV and SARS-CoV-2 have been found circulating among wild animals and as a result present a threat to our public health security. Thus, GS-5245 may not only prove to be important for treating SARS-CoV-2 infections but could also be deployed to outbreaks of new coronaviruses that may emerge in the future. Despite the high degree of conservation in the RdRp and M^pro^, coronaviruses can evolve to escape single-agent therapies in a laboratory setting (*41, 42*). Like HIV-1 antiviral therapy treatments that combine antivirals with different mechanisms or action, such as protease and polymerase inhibitors (*43*), we demonstrate that combination oral antiviral therapy to treat acute SARS-CoV-2 infections may be advantageous to increase efficacy of antiviral interventions. An additional benefit of combining oral antiviral therapies to treat COVID-19 may be the reduced risk of monotherapy drug-resistant SARS-CoV-2 variants as has been documented in remdesivir-treated immunocompromised patients (*44–46*). Moreover, in vitro selection studies demonstrate that SARS-CoV-2 can acquire mutations that confer the Mpro protease with resistance to nirmatrelvir (*47*). Therefore, it will be critical to explore combination therapy in humans to not only increase the efficacy of antivirals that have different viral targets but also to potentially reduce risk of antiviral resistance in the setting of monotherapy, particularly in immunocompromised patients who invariably experience prolonged viral replication. Combination antiviral interventions may also diminish COVID-19 progression to Post-Acute Sequelae of SARS-CoV-2 infection (PASC)/long COVID manifestations (*48*). Moreover, as SARS-CoV-2 infectious virus rebound following Paxlovid™ treatment occurs in patients (*49, 50*), combination drug intervention strategies may be clinically useful in reducing or eliminating cases of viral rebound.

Altogether, our data indicates that GS-5245 may have clinical utility against highly transmissible SARS-CoV-2 variants as well as zoonotic coronaviruses that may emerge in the future. Moreover, GS-5245 demonstrated efficacy against common-cold human alphacoronavirus NL63 and pre-emergent SARS-CoV-related viruses, underlining the broad antiviral efficacy of this drug and its potential treatment of common-cold human coronaviruses, which can progress to severe life-threatening illnesses in the very young and aged (*51*). GS-5245 is currently being investigated in two international Phase 3 trials in standard and high risk COVID-19 patients (ClinicalTrials.gov: NCT05715528 and NCT05603143). The present in vitro and in vivo data support the continued clinical evaluation of GS-5245 for the treatment of COVID-19 and potentially other coronaviruses.

### Limitations of the study

Despite the strong efficacy of GS-5245 in vivo, our study has limitations. The breadth of protection was only tested in mice and not in additional animal models of coronavirus pathogenesis. While remdesivir exhibits clinical efficacy across broad patient populations and stages of COVID-19 disease, the testing of GS-5245’s clinical efficacy is ongoing. The clinical efficacy of GS-5245 may be established based on ongoing controlled, randomized, and powered Phase III clinical trials in humans; the study results are pending. Similarly, while GS-5245 and nirmatrelvir exhibited enhanced efficacy when tested in combination in SARS-CoV-2-infected mice, the clinical efficacy of GS-5245 in combination with Paxlovid™ based on properly designed human clinical studies has not yet been established. Therefore, it will be important to continue evaluating both GS-5245 monotherapy and combination therapy with Paxlovid™ in future human clinical trials.

## ACKNOWLEDGEMENTS

This work was supported by an NIAID AVIDD U19AI171292 to R.S.B, R01 AI132178 and R01AI132178-04S1 to T.P.S. and R.S.B., and an NIH animal models contract (HHSN272201700036I) to R.S.B. Histopathology was performed by the Pathology Services Core at the University of North Carolina-Chapel Hill, which is supported in part by an NCI Center Core Support Grant (5P30CA016080-42). D.R.M. is funded by a Hanna H. Gray Fellowship from the Howard Hughes Medical Institute.

## AUTHOR CONTRIBUTIONS

Conceived the study: D.R.M., A.S., and T.P.S.; designed experiments: D.R.M., J.Y.F., J.P.B. T.C., R.L.M., R.S.B., K.T.B., R.B., A.S., and T.P.S.; performed laboratory experiments: D.R.M., F.R.M., M.R.Z., K.G., G.D.l.C., J.J.W., A.J.B., W.S.C., S.R.M., S.D., S.M., A.S., and T.P.S.; provided critical reagents: D.B., K.B., A.N., R.K., A.N., R.B, J.P.B., J.F., T.C., R.L.M.; wrote the first draft of the paper: D.R.M. edited the manuscript: D.R.M., D.B., K.T.B., R.B, J.P.B., J.P., J.F., T.C., R.L.M.,R.S.B, A.S., and T.P.S.; all authors reviewed and approved the manuscript.

## STATISTICAL ANALYSIS

The statistical analysis performed using SAS Software version 9.4 (SAS Institute Inc., Cary, NC, USA) and the R statistical package version 3.6.1 (Vienna, Austria). Statistical tests used are described in the figure legends.

## MATERIALS AND METHODS

### Viruses

Recombinant viruses utilized for in vitro studies include SARS-CoV Urbani expressing nanoluciferase (SARS-nLuc, nLuc replaces ORF7) and MERS-CoV EMC Strain, SARS-CoV-2 WA1, Bat SARS-related CoV RsSHC014, HCoV-NL63 were created from molecular cDNA clones as described (*52–54*) previously. SARS-CoV-2 Omicron BA.1 isolate was obtained as a gift from Dr. Yoshihiro Kawaoka from the University of Wisconsin, Madison.

### Preclinical experiments

#### Mouse strains and infections

Eight-week-old female BALB/c mice were purchased from Envigo (#047). Eight-week-old female K18-humanized ACE2 mice on a B6 background were purchased from Jackson Laboratory (#034860). 288/330^++^ dipeptidyl peptidase 4 (DPP4) mice were bred at the University of North Carolina at Chapel Hill and were described previously (*34*). For SARS-CoV-2 infections in BALB/c mice, a mouse-adapted SARS-CoV-2 virus (MA10) was used and mice were infected with 1 x 10^4^ PFU intranasally (*28*). For SARS-CoV infections in BALB/c mice, an intranasal infectious of 1 x 10^4^ PFU was used (*32*). For RsSHC014-CoV and SARS-CoV-2 Omicron (BA.1) infections, K18-hACE2 mice were infected with 1 x 10^4^ PFU intranasally. Finally, 288/330^++^ DPP4 mice were infected intranasally with 5 x 10^4^ PFU with a mouse-adapted MERS-CoV (maM35c4) virus as described previously (*33, 34*). Infected mice were weighed daily and were monitored for signs of disease in all infection and treatment studies.

#### GS-5245, molnupiravir, and PF-07321332 formulations

GS-5245 was synthesized at Gilead Sciences, Inc., and their chemical composition and purity was quality controlled by nuclear magnetic resonance, high resolution mass spectrometry, and high-performance liquid chromatography. Molnupiravir was purchased from MedChemExpress (Monmouth Junction, NJ). Nirmatrelvir (PF-07321332) was purchased from MedChemExpress (Monmouth Junction, NJ) or WuXi AppTec (Shanghai, China). GS-5245 was solubilized in 2.5% dimethyl sulfoxide, 10% Kolliphor HS-15; 10% Labrasol; 2.5% propylene glycol; 75% water; pH 2-3 or 10% ethanol, 90% propylene glycol for mouse *in vivo* studies. Molnupiravir was solubilized in 2.5% Kolliphor RH 40, 10% polyethylene glycol, 87.5% water for mouse *in vivo* studies. PF-07321332 was formulated in 10% ethanol, 90% propylene glycol. Oral antiviral drugs were made available to UNC Chapel Hill under an existing material transfer agreement with Gilead Sciences, Inc.

#### Structural analysis of nsp12 conservation

A model of the pre-incorporation state of the GS-5245 active metabolite in the SARS-CoV-2 polymerase complex was based on the cryo-EM structure 6XEZ (*55*) and previously described (*56*). Homology models using Schrödinger Release 2022-2 (Prime, Schrödinger, LLC, New York, NY) were generated for the nsp12 subunit were generated for the viral strains used in this study.

#### Mouse GS-5245 pharmacokinetic studies

Mouse pharmacokinetic (PK) studies were performed at LabCorp Drug Development. Briefly, n=4 female BALB/c and n=3 female and n=3 male mice from each different strain of K18-hACE2 and 288/330^++^ DPP4 were dosed with 30 mg/kg of GS-5245 by oral gavage. Plasma concentrations (µM) of GS-411524 were quantitated in each mouse strain at select timepoints over 24 hours as described previously (*27*).

#### Antiviral activity of GS-5245 against SARS-CoV-2

We employed an antiviral assay as described (*27*). Briefly, human lung epithelial cell line A549 (ATCC # CCL185) stably expressing hACE2 were plated at a density of 20,000 cells per well in 100 µL in black-walled clear-bottom 96-well plates 24 hours prior to infection. GS-5245, RDV, GS-441524, remdesivir, and PF-07321332 were diluted in 100% DMSO (1:2) resulting in a 1000X dose response from 10 to 0.039 mM 10 to 0.039 µM. All conditions were performed in triplicate. In a Biosafety Level 3 Laboratory (BSL3), medium was removed, and cells were infected with 100 µL SARS-CoV-2 nLUC (multiplicity of infection (MOI) = 0.5) for 1 hour at 37 °C. After this incubation, virus was removed and wells were washed (150 µL) with infection media (Dulbecco’s Modified Eagle’s Medium (DMEM), 4% fetal bovine serum (FBS), 1X antibiotic/antimycotic). Infection media (100 µL) containing diluted drug in dose response was then added. DMSO remained at 0.1% across the dilution series. Plates were incubated at 37 °C for 48 hours. NanoGlo assay was performed at 48 hpi. Sister plates were exposed to drug but not infected to gauge cytotoxicity using a CellTiter-Glo assay (CTG, Promega) at 48 hours post treatment. Values were normalized to the uninfected and infected DMSO controls (0% and 100% infection, respectively). Data was fit using a four-parameter non-linear regression analysis using GraphPad Prism. EC_50_ and CC_50_ values were then determined as the concentration reducing the signal by 50%.

#### Antiviral activity of GS-5245 against HCoV-NL63

An antiviral assay was performed with recombinant NL63 reporter virus expressing nanoluciferase as described (*57*). Briefly, LLCMK2 cells stably expressing TMPRSS2 were seeded at 20,000 cells per well and infected with NL63 nLUC at an MOI of 0.02 for 1 hr after which virus was removed, cultures were washed with medium, and a dose response of each drug was added in triplicate. After 48 hr, virus replication was quantitated by measuring nanoluciferase expression. For cytotoxicity, sister plates were exposed to the same dose response as those infected. Cytoxicity was quantitated using CellTiter Glo assay at 48 hours post exposure. Three independent studies were performed. Values were normalized to the uninfected and infected DMSO controls (0% and 100% infection, respectively). Data was fit using a four-parameter non-linear regression analysis using GraphPad Prism. EC_50_ and CC_50_ values were then determined as the concentration reducing the signal by 50%.

#### Antiviral activity in primary human airway epithelial cells

Human airway epithelial (HAE) cell cultures were obtained from the Tissue Procurement and Cell Culture Core Laboratory in the Marsico Lung Institute/Cystic Fibrosis Research Center at UNC. Prior to infection, HAE were washed with PBS and moved into ALI media containing a dose response of 10, 1, or 0.1µM GS-5245 or DMSO. HAE were infected at an MOI of 0.5 for 1.5 hr at 37°C after which virus was removed, cultures were washed with PBS and then incubated at 37°C for 72hr. Virus replication/titration was performed as previously described. Similar data was obtained in two independent studies using cells from two different patient donors. Cytotoxicity was measured via ToxiLight Assay (Lonza) in triplicate HAE cell cultures treated with 10, 1, or 0.1µM GS-5245 or DMSO. DMSO remained at 0.1% for all conditions. As a positive control, duplicate wells were exposed to Promega nanoluciferase lysis buffer for 10 minutes prior to Toxilight Assay.

#### Animal care

Animal efficacy studies were performed in accordance with the recommendations for care and use of animals by the Office of Laboratory Animal Welfare (OLAW), National Institutes of Health and the Institutional Animal Care and Use Committee (IACUC) protocol number: 20-059 at University of North Carolina (UNC permit no. A-3410-01). All mice were anesthetized prior to viral inoculation and great efforts were undertaken to reduce animal suffering. Mice were fed standard chow diets and housed in groups of five.

#### Lung pathology scoring

Acute lung injury was quantified with two distinct lung pathology scoring tools. The first scoring tool is the Matute-Bello scoring system developed by the American Thoracic Society (ATS), and the second scoring tool is the Diffuse Alveolar Damage (DAD) to score lung damage caused by acute viral infections as described previously (*14*). Lung scoring and scoring analyses were performed by a Board-Certified Veterinary Pathologist who was blinded to the treatment groups. Lung pathology slides were read and scored at 600X total magnification.

#### Laboratory biosafety

Cell culture and animal studies were approved by the UNC Institutional Biosafety Committee approved under laboratory and animal protocols used in the Baric laboratory. All work described here, including coronavirus work, was performed with approved standard operating procedures for SARS-CoV, SARS-CoV-2, and MERS-CoV in a biosafety level 3 (BSL-3) facility which met and exceeded requirements recommended in the Microbiological and Biomedical Laboratories, by the U.S. Department of Health and Human Service, the U.S. Public Health Service, and the U.S. Center for Disease Control and Prevention (CDC), and the National Institutes of Health (NIH).

## Quantification and statistical analysis

Statistical analyses were performed in Prism version 9.0, SAS Software version 9.4, and the R statistical package version 3.6.1. Figure legends describe each statistical test used in each figure.

## Materials availability

Material and reagents generated in this study will be made available upon installment of a standard material transfer agreement (MTA) through UNC, while other reagents and viruses are available through BEI.

## Supplementary Materials

**Fig. S1.**
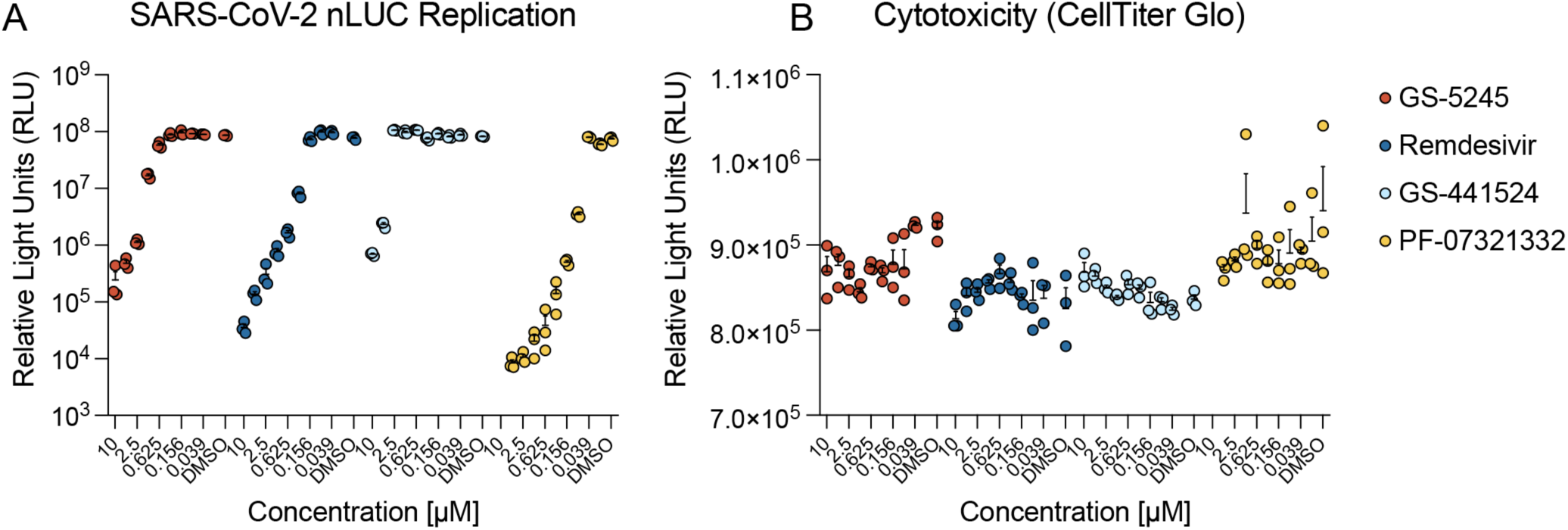
Measuring the antiviral and cytotoxic activity of GS-5245, GS-441524, remdesivir, and PF-07321332 in A549-hACE2 cells (related to Fig. 1). (A) A549-hACE2 were plated at 20,000 cells per well and infected with SARS-CoV-2 nLUC at an MOI of 0.5 for 1hr after which virus was removed, cultures were washed with medium and then a dose response of each drug was added in triplicate. After 48hr, virus replication was quantitated by measuring nLUC expression. (B) For cytotoxicity, sister plates were exposed to the same dose response as those infected. Cytotoxicity was quantitated using CellTiter Glo assay at 48 hr post exposure.Fig. S2. Generation, recovery, and characterization of an NL63 infectious clone (ic) expressing nanoluciferase (nLuc) (related to Fig. 1).

**Fig. S2.**
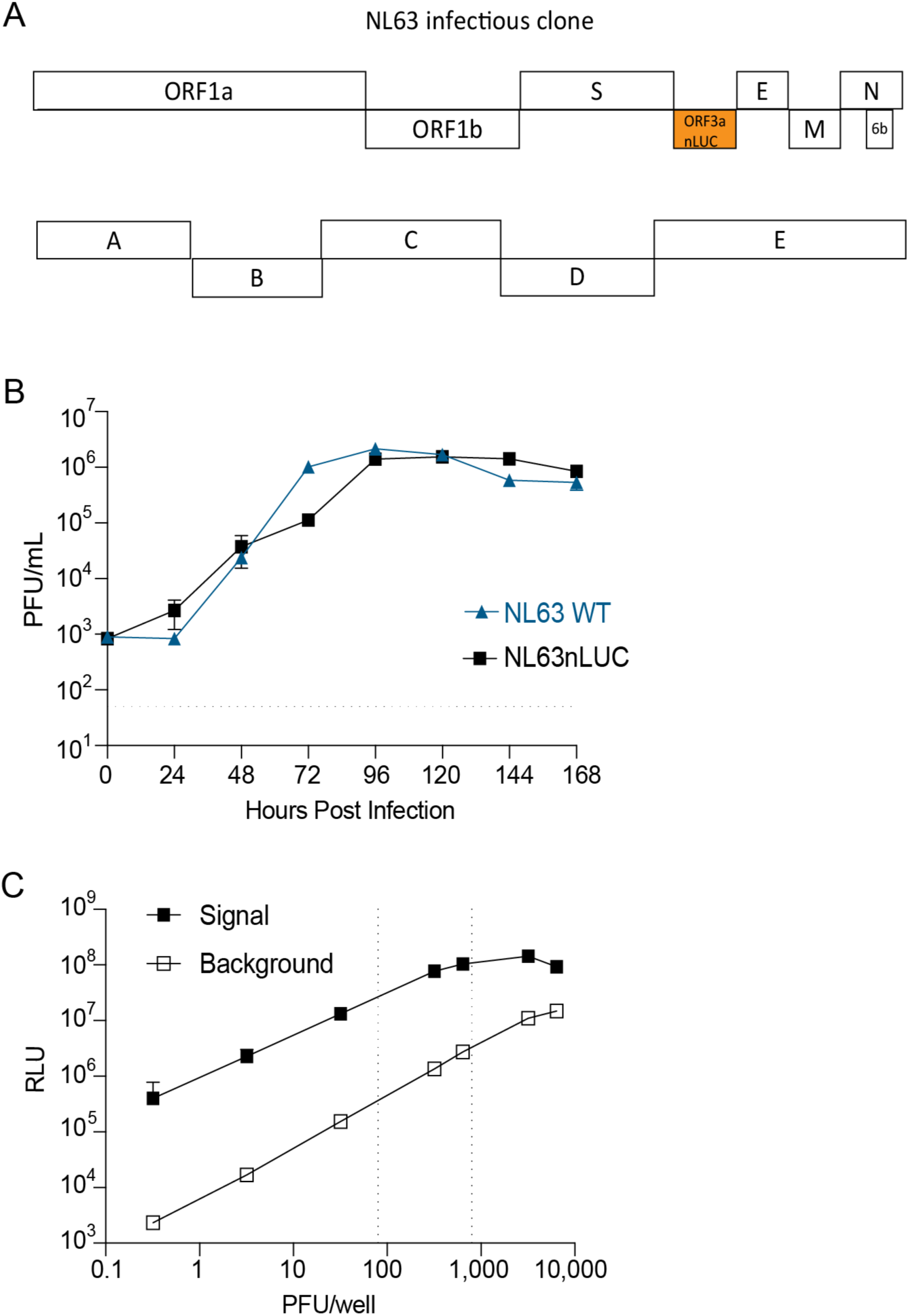
Generation, recovery, and characterization of an NL63 infectious clone (ic) expressing nanoluciferase (nLuc) (related to Fig. 1). (A) Infectious clone diagram of NL63 open reading frames (top) and the fragments used in assembly (bottom). ORF3a was replaced with a gene encoding nanoluciferase. (B) Growth kinetics of icNL63 (blue, triangle) and icNL63-nLuc (black, square). (C) Luciferase signal of icNL63-nLuc 72 hpi. Background luciferase in virus without infecting cells (open square). Luciferase signal after infecting cells (filled square).

**Fig. S3.**
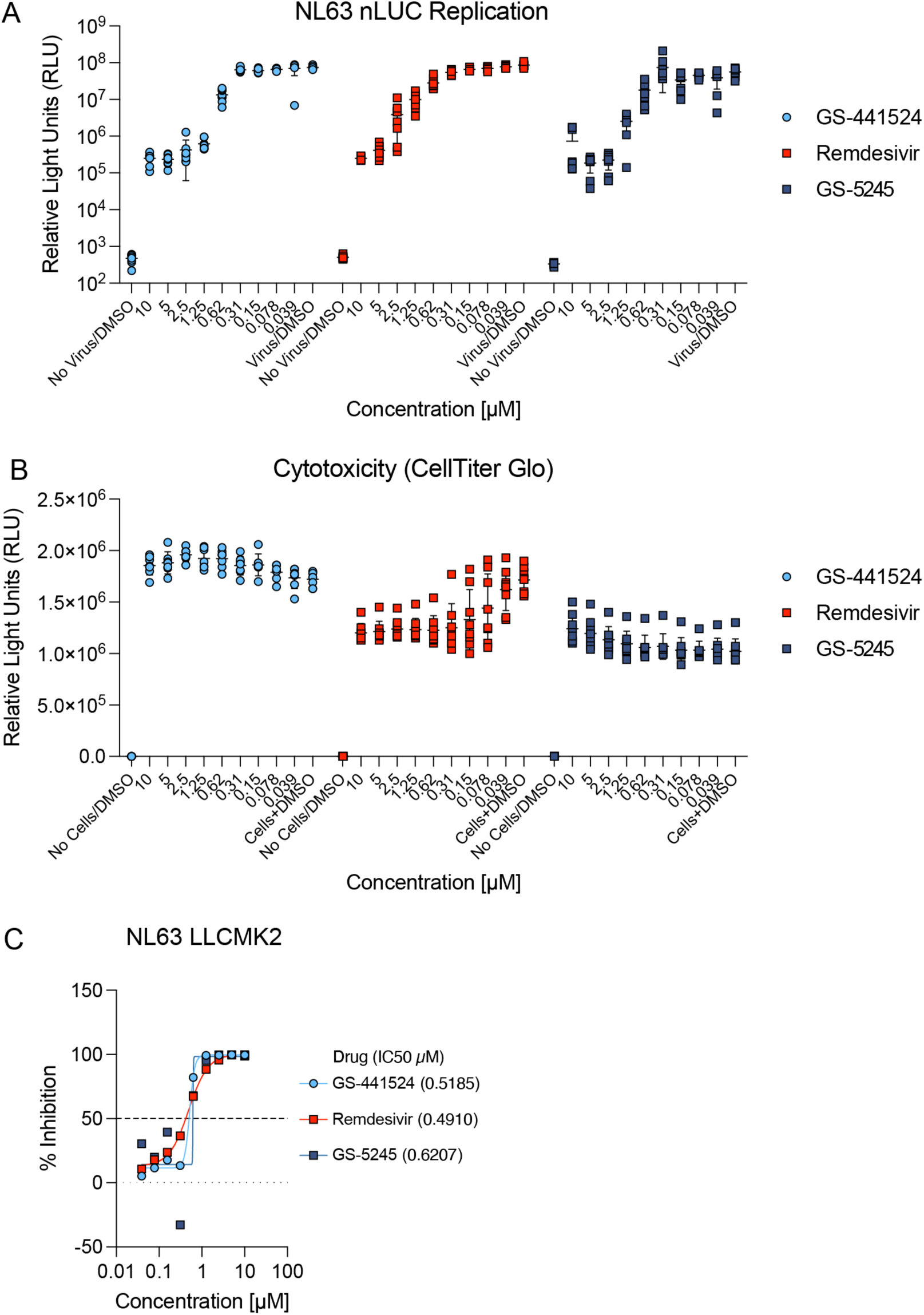
The antiviral activity of GS-5245 against human coronavirus NL63 (related to Fig. 1). (A) LLCMK2 cells stably expressing TMPRSS2 were seeded at 20,000 cells per well and infected with NL63 nLUC at an MOI of 0.02 for 1 hr after which virus was removed, cultures were washed with medium and then a dose response of each drug was added in triplicate. After 48 hr, virus replication was quantitated by measuring nLUC expression. (B) For cytotoxicity, sister plates were exposed to the same dose response as those infected. Cytotoxicity was quantitated using CellTiter Glo assay at 48 hr post exposure. (C) The percent antiviral inhibition for each drug is shown.

**Fig. S4.**
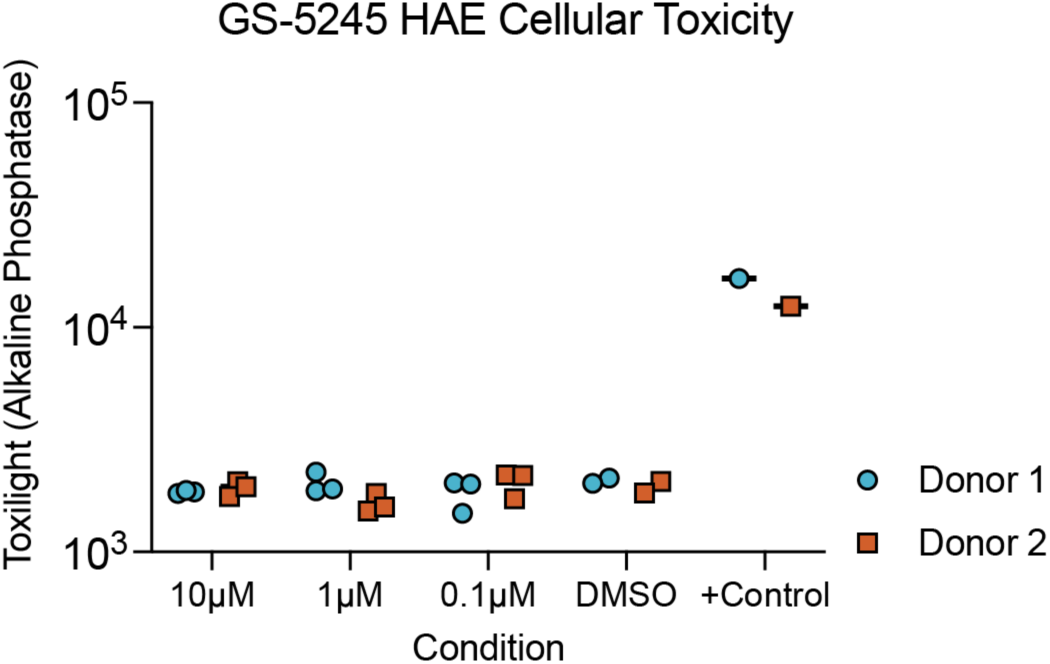
Cytotoxicity of GS-5245 in HAE (related to Fig. 1). Primary human airway epithelial cell cultures from donors 1 and 2 were treated with different doses of GS-5245 or DMSO for 72hr in triplicate, after which cytotoxicity was assessed by ToxiLight Assay which measures alkaline phosphatase released by dead and dying cells. As a positive control (+ Control), duplicate wells were exposed to Promega nanoluciferase lysis buffer for 10 minutes prior to Toxilight Assay.

**Fig. S5.**
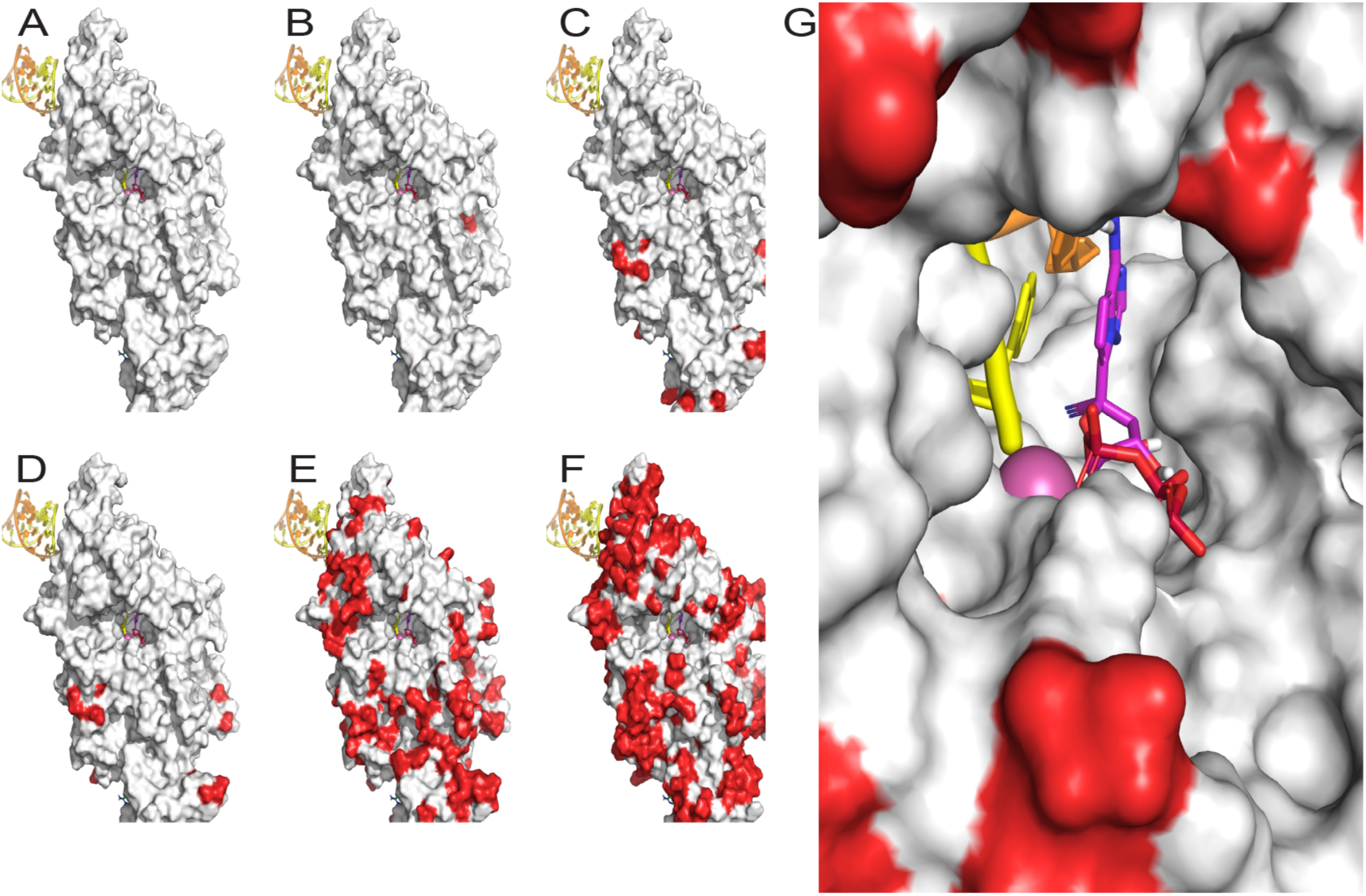
Homology models of nsp12 show the polymerase active site is well conserved across coronaviruses (related to Fig. 1). (A) Model of the pre-incorporated GS-5245 active triphosphate metabolite (purple) in SARS-CoV-2 WA-1 nsp12 was based on the cryo-EM structure 6XEZ. (B) A homology model of Omicron BA.1 with the single mutation, P323L (shown in red), was generated from the WA-1 model. Additional homology models were generated for (C) SARS-CoV, (D) RsSHC014-CoV, (E) MERS-CoV EMC strain and (F) HCoV-NL63, where non-conserved residues with respect to WA-1 are also shown in red. (G) A detail of the NL63 active site, the least conserved virus among this set, shows the active site, and particularly the residues which recognize the important 1’-CN of the active triphosphate produced from GS-5245, are completely conserved.

**Fig. S6.**
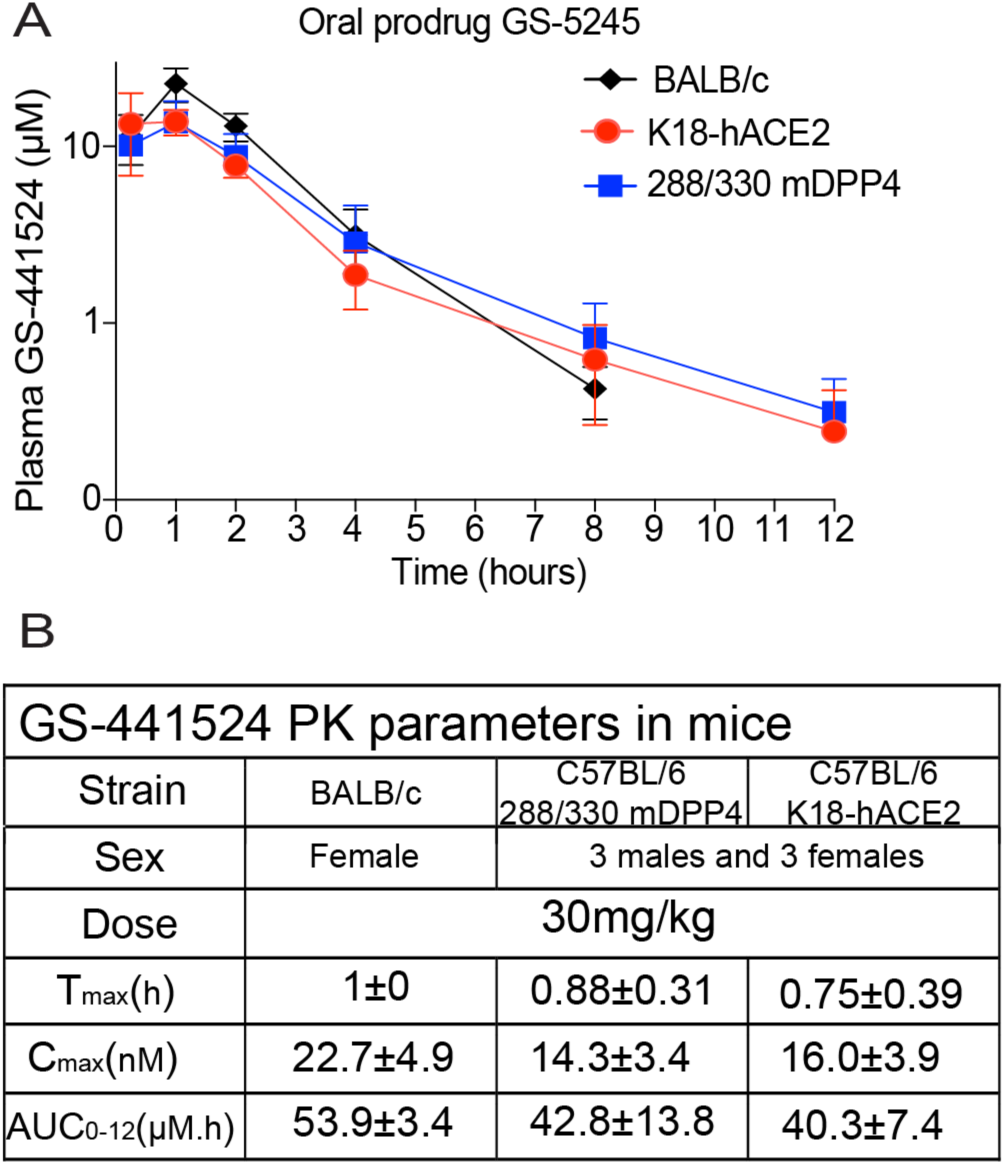
Pharmacokinetics of GS-441524 following oral administration of GS-5245 in different mouse strains (related to Figs. 2, 3, 4, and 5) (A) Plasma pharmacokinetics of GS-441524 in different mouse strains. (B) BALB/c, K18-hACE2, and 288/330^++^ DPP4 mice were orally gavaged with 30 mg/kg of GS-5245 and plasma concentrations (µM) of GS-411524 were quantified. A total of n=4 females in BALB/c and n=3 males and n=3 females per mouse strain in K18-hACE2, and 288/330^++^ DPP4 were used.

**Fig. S7.**
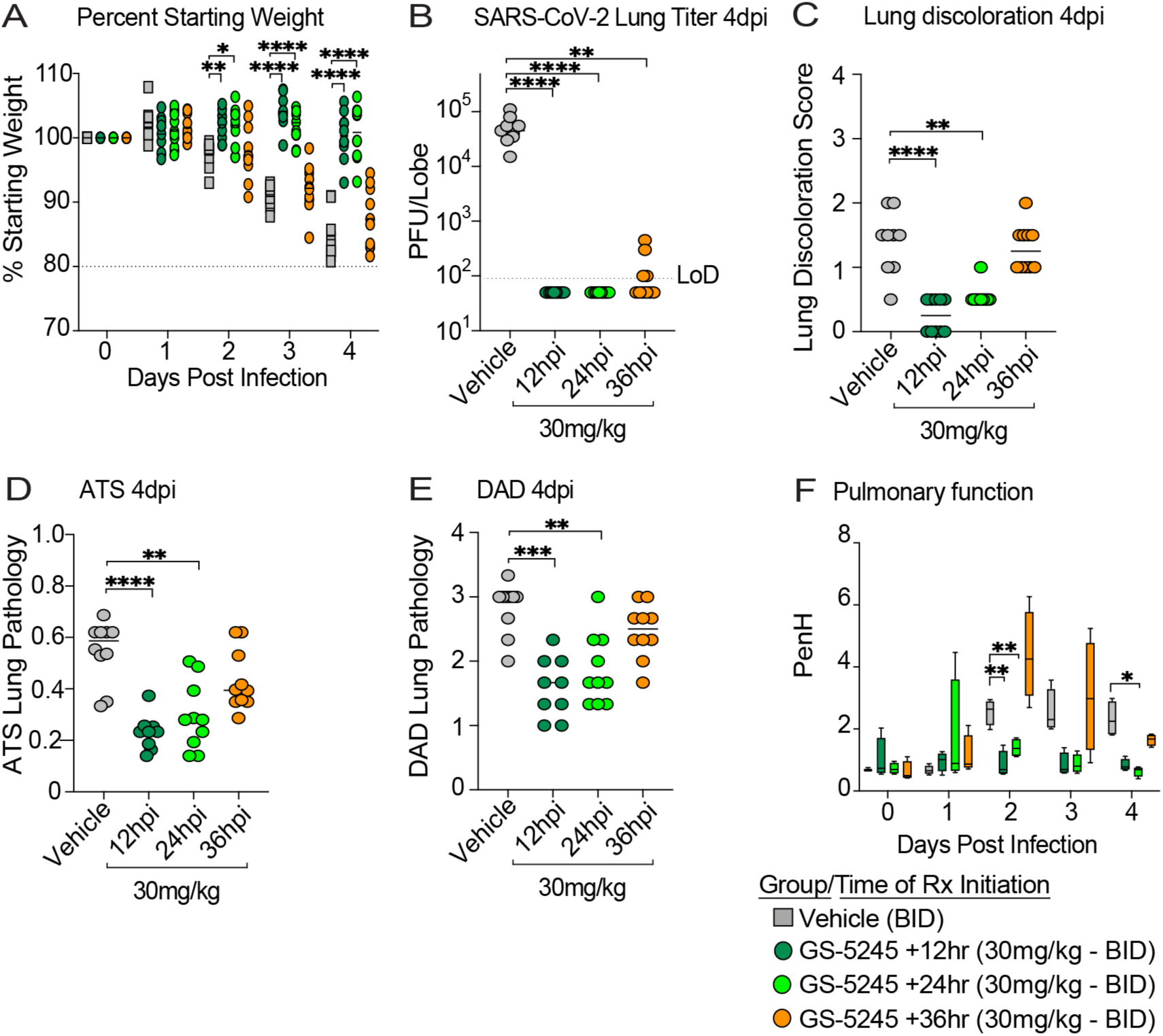
30 mg/kg GS-5245 treatment kinetics in SARS-CoV-2-infected BALB/c mice (related to Fig. 2). (A) Percent starting weight through 4 dpi in 10-week-old female BALB/c mice infected with SARS-CoV-2 MA10 at 1 x 10^4^ PFU. Mice were treated BID with vehicle (gray squares), 30 mg/kg GS-5245 at 12 hpi (dark green circles), 30 mg/kg GS-5245 at 24 hpi (light green circles), and 30 mg/kg GS-5245 at 36 hpi (orange circles). N=10 mice per group. Asterisks denote p values from a two-way ANOVA after a Dunnett’s multiple comparisons test. * p ≤ 0.05, ** p ≤ 0.01, *** p ≤ 0.001, **** p ≤ 0.0001 (B) SARS-CoV-2 MA10 lung infectious viral titers 4 dpi in mice treated with vehicle or 30 mg/kg GS-5245 at 12, 24, and 36 hpi. Asterisks denote p values from a Kruskal-Wallis test after a Dunnett’s multiple comparisons test. (C) Macroscopic lung discoloration at 4 dpi in therapeutically treated mice with GS-5245 at various intervention timepoints compared to vehicle. Asterisks denote p values from a Kruskal-Wallis test after a Dunnett’s multiple comparisons test. (D) Microscopic ATS acute lung injury pathology scoring at 4 dpi in vehicle vs. GS-5245-treated mice. Asterisks denote p values from a Kruskal-Wallis test after a Dunnett’s multiple comparisons test. (E) Microscopic DAD acute lung injury pathology scoring at 4 dpi in vehicle vs. GS-5245-treated mice. Asterisks denote p values from a Kruskal-Wallis test after a Dunnett’s multiple comparisons test. (F) Pulmonary function monitored by whole-body plethysmography from day zero through 4 dpi in SARS-CoV-2-infected 30 mg/kg GS-5245 vs. vehicle treated mice. Asterisks denote p values from a two-way ANOVA after a Dunnett’s multiple comparisons test.

**Fig. S8.**
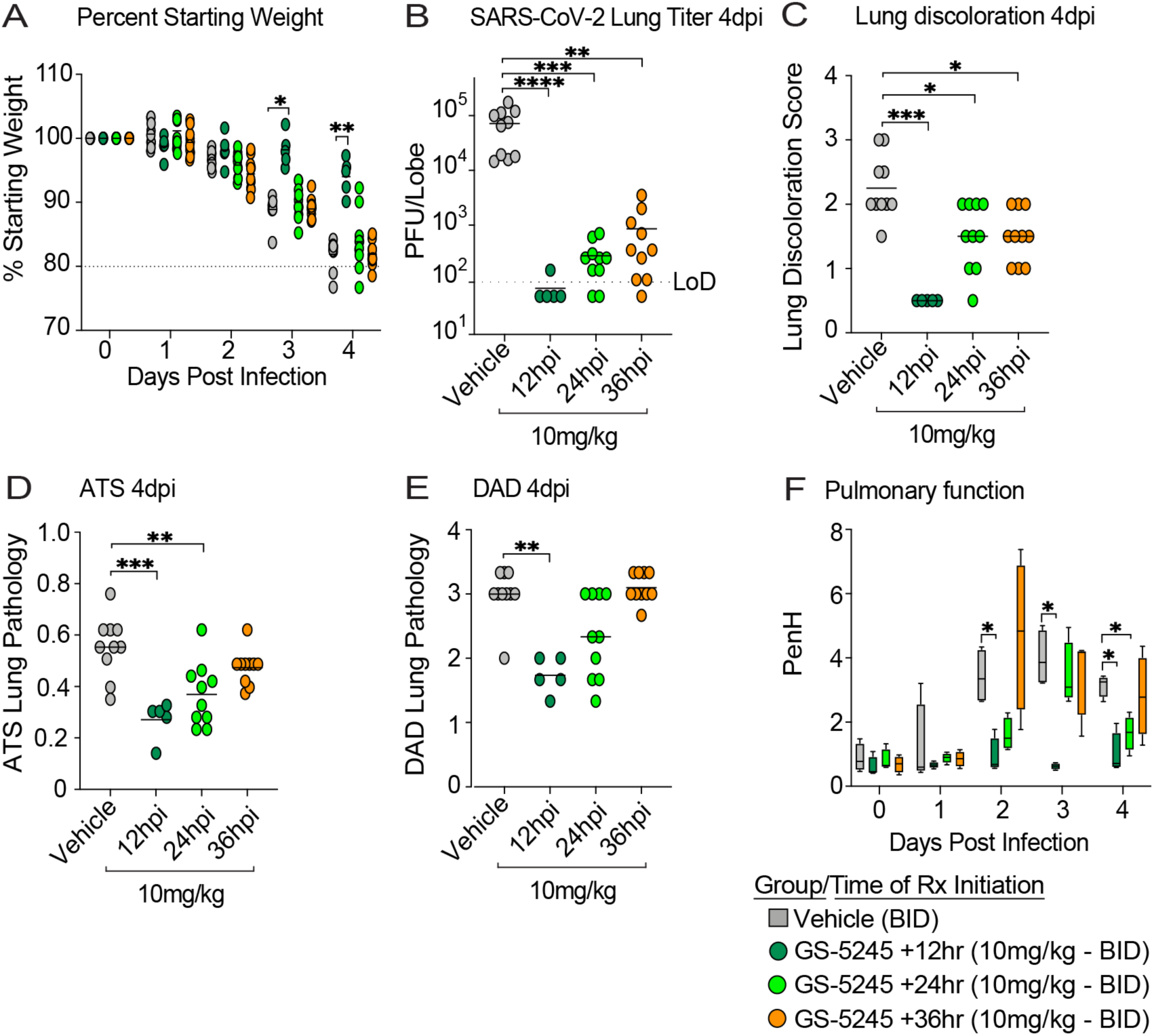
10 mg/kg GS-5245 treatment kinetics in SARS-CoV-2-infected BALB/c mice (related to Fig. 2) (A) Percent starting weight through 4 dpi in 10-week-old female BALB/c mice infected with SARS-CoV-2 MA10 at 1 x 10^4^ PFU. Mice were treated BID with vehicle (gray squares), 10 mg/kg GS-5245 at 12 hpi (dark green circles), 10 mg/kg GS-5245 at 24 hpi (light green circles), and 10 mg/kg GS-5245 at 36 hpi (orange circles). N=10 mice per group. Asterisks denote p values from a two-way ANOVA after a Dunnett’s multiple comparisons test. * p ≤ 0.05, ** p ≤ 0.01, *** p ≤ 0.001, **** p ≤ 0.0001 (B) SARS-CoV-2 MA10 lung infectious viral titers 4 dpi in mice treated with vehicle or 10 mg/kg GS-5245 at 12, 24, and 36 hpi. Asterisks denote p values from a Kruskal-Wallis test after a Dunnett’s multiple comparisons test. (C) Macroscopic lung discoloration at 4 dpi in therapeutically treated mice with GS-5245 at various intervention timepoints compared to vehicle. Asterisks denote p values from a Kruskal-Wallis test after a Dunnett’s multiple comparisons test. (D) Microscopic ATS acute lung injury pathology scoring at 4 dpi in vehicle vs. GS-5245-treated mice. Asterisks denote p values from a Kruskal-Wallis test after a Dunnett’s multiple comparisons test. (E) Microscopic DAD acute lung injury pathology scoring at 4 dpi in vehicle vs. GS-5245-treated mice. Asterisks denote p values from a Kruskal-Wallis test after a Dunnett’s multiple comparisons test. (F) Pulmonary function monitored by whole-body plethysmography from day zero through 4 dpi in SARS-CoV-2-infected 10 mg/kg GS-5245 vs vehicle treated mice. Asterisks denote p values from a two-way ANOVA after a Dunnett’s multiple comparisons test.

